# Meiotic chromosome synapsis depends on multivalent SYCE1-SIX6OS1 interactions that are disrupted in cases of human infertility

**DOI:** 10.1101/2020.02.04.934372

**Authors:** Fernando Sánchez-Sáez, Laura Gómez-H, Orla M. Dunne, Cristina Gallego-Páramo, Natalia Felipe-Medina, Manuel Sánchez-Martín, Elena Llano, Alberto M. Pendas, Owen R. Davies

## Abstract

Meiotic reductional division is dependent on the synaptonemal complex (SC), a supramolecular protein assembly that mediates homologous chromosomes synapsis and promotes crossover formation. The mammalian SC is formed of eight structural components, including SYCE1, the only central element protein with known causative mutations in human infertility. We combine mouse genetics, cellular and biochemical studies to reveal that SYCE1 undergoes multivalent interactions with SC component SIX6OS1. The N-terminus of SIX6OS1 binds and disrupts SYCE1’s core dimeric structure to form a 1:1 complex, whilst their downstream sequences provide a distinct second interface. These interfaces are separately disrupted by SYCE1 mutations associated with non-obstructive azoospermia and premature ovarian failure, respectively. Mice harbouring SYCE1’s POF mutation and a targeted deletion within SIX6OS1’s N-terminus are infertile with failure of chromosome synapsis. We conclude that both SYCE1-SIX6OS1 binding interfaces are essential for SC assembly, thus explaining how SYCE1’s reported clinical mutations give rise to human infertility.

## Introduction

Meiotic cell division is defined by a unique and highly dynamic programme of events that result in homologous chromosome synapsis, crossover formation and subsequent homologue segregation into haploid germ cells (*1–3*). In mammals, homologous chromosome pairs are established through inter-homologue recombination searches from up to 400 induced double-strand breaks (DBSs) per cell (*4*). Once established, local recombination-mediated alignments are converted into the single continuous synapsis of aligned homologous chromosomes through the zipper-like assembly of the synaptonemal complex (SC) (*5*). The SC’s supramolecular protein structure mediates continuous 100-nm tethering between homologous chromosome axes and provides the necessary three-dimensional framework for crossover formation (*2*). Following SC disassembly, crossovers provide the sole physical links between homologues at metaphase I, so are essential for ensuring correct homologue segregation in addition to providing genetic diversity (*2*).

The SC has an iconic and highly conserved tripartite structure that has been observed across meiotically-reproducing eukaryotes (*6*). This consists of lateral elements (LEs) that coat the two homologous chromosome axes and a midline central element (CE), with a series of transverse filaments that bind together these longitudinal electron-dense structures (Figure 1a) (*7*). The protein components of the mammalian SC have been identified as transverse filaments protein SYCP1 (*8*), CE proteins SYCE1-3, SIX6OS1 and TEX12 (*9–12*), and LE proteins SYCP2 and SYCP3 (*13, 14*). All transverse filament and CE components are essential for SC assembly, and their individual disruption leads to infertility owing to meiotic arrest with failure of DSB repair (*10, 11, 15-18*). In contrast, disruption of LE components produces a sexual dimorphism of male infertility and female subfertility (*19, 20*), with SYCP3 deficiency in females promoting germ cell aneuploidy and embryonic death (*21*).

**Figure 1.**
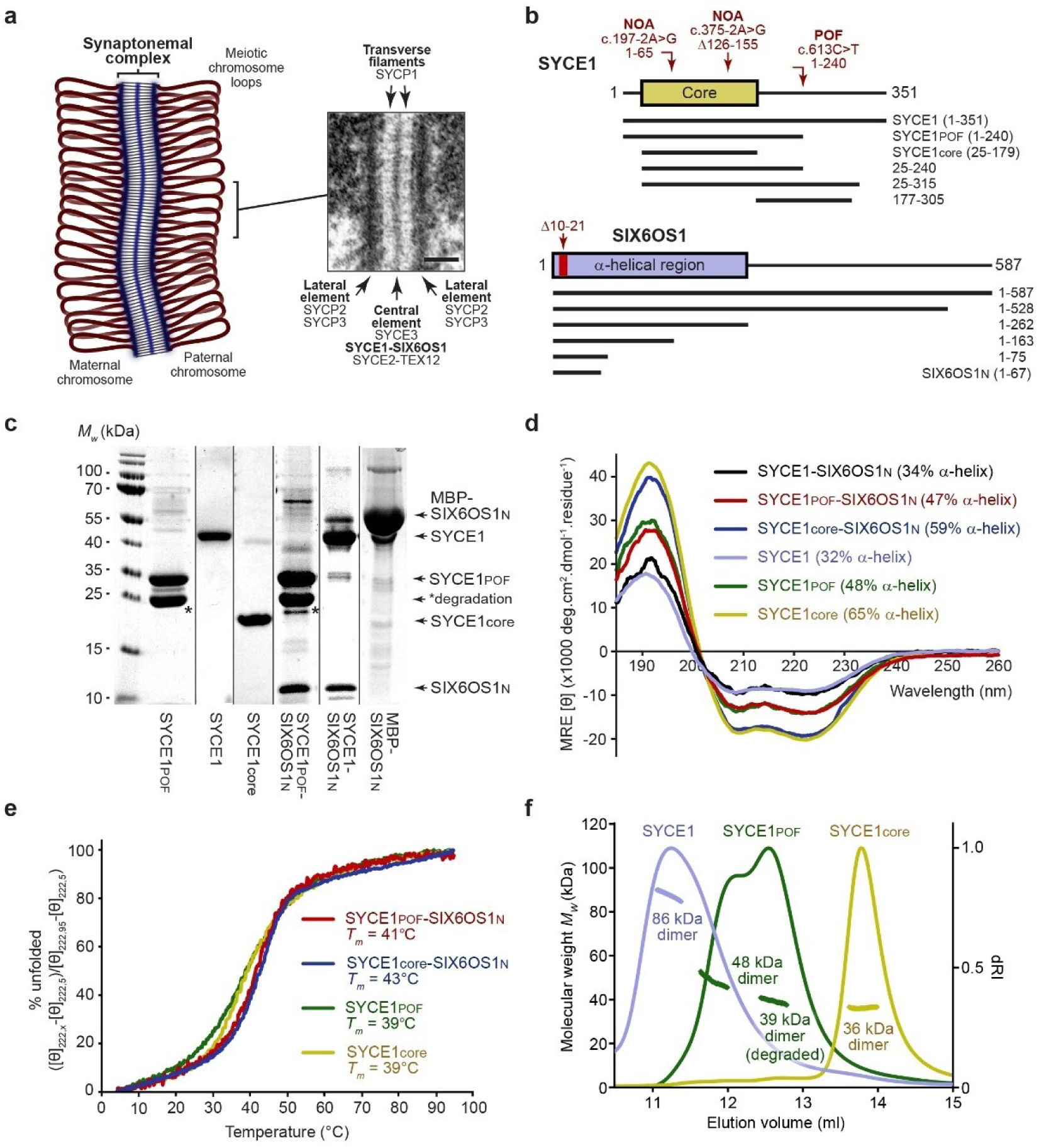
SYCE1POF retains its core dimeric structure. (**a**) Schematic of the synaptonemal complex (SC) binding together a pair of homologous chromosomes in meiosis, modified from Dunne and Davies (*28*), with inset SC electron micrograph reproduced from Kouznetsova, Benavente, Pastink and Hoog (*30*). Scale bar, 100 nm. The SC has a tripartite structure of three electron-dense longitudinal structures - chromosome-bound LEs and a midline central element – that are connected together by transverse filaments. Protein components of the mammalian SC are indicated. (**b**) Human SYCE1 (top) and SIX6OS1 (bottom) sequence schematics indicating the location and consequence of infertility-associated mutations of SYCE1 and Δ10-21 internal deletion of SIX6OS1, alongside the principal constructs used in this study. (**c**) SDS-PAGE analysis of the purified recombinant proteins used in this study; *indicates the dominant degradation product of SYCE1POF of apparent molecular weight consistent with C-terminal truncation to the 25-179 core. (**d**) Far UV circular dichroism (CD) spectra recorded between 260 nm and 185 nm in mean residue ellipticity, MRE ([θ]) (x1000 deg.cm^2^.dmol^−1^.residue^−1^). Data were deconvoluted using the CDSSTR algorithm revealing the helical contents indicated. (**e**) CD thermal denaturation recording the CD helical signature at 222 nm between 5°C and 95°C, as % unfolded; estimated melting temperatures (*Tm*) are indicated. (**f**) Size exclusion chromatography-multi angle light scattering (SEC-MALS) analysis; light scattering (LS) and differential refractive index (dRI) are shown as solid and dashed lines respectively, with fitted molecular weights (Mw) plotted as diamonds across elution peaks. SYCE1core (yellow), SYCE1POF (green) and full-length SYCE1 (blue) are dimeric species of 36 kDa, 48 kDa (39 kDa for the degradation product) and 86 kDa, respectively (theoretical dimers – 37 kDa, 55 kDa and 80 kDa). Data for SYCE1core and full-length SYCE1 are reproduced from Dunne and Davies (*28*).

In recent years, a variety of cellular imaging, biochemical and structural biology approaches have begun to uncover the molecular structures, interactions and mechanisms responsible for mammalian SC assembly. SYCP1 self-assembles into a supramolecular lattice that provides the underlying 100-nm synapsis between chromosome axes (*22, 23*), whilst SYCP3 assembles into regularly-repeating filaments that support chromosomal looping (*24, 25*). The five CE proteins provide essential structural supports for the SYCP1 lattice that enable its continuous and cooperative extension along the entire chromosome length. In this capacity, CE proteins have been categorised as synaptic initiation factors (SYCE3, SYCE1 and SIX6OS1) and elongation factors (SYCE2 and TEX12), of which their disruption leads to complete loss of tripartite SC structure and failure of extension of short SC-like stretches, respectively (*10, 11, 16-18*). Of synaptic initiation factors, SYCE3 forms dimers that undergo potentially limitless self-assembly (*26, 27*), SYCE1 forms anti-parallel dimeric assemblies (*28*), and SIX6OS1 is an SYCE1-interacting protein of unknown structure (*11*). These likely act as short-range structural supports between SYCP1 molecules, possibly in transverse, longitudinal and vertical orientations to stabilise a local three-dimensional SYCP1 lattice (*22*). In contrast, SYCE2 and TEX12 exist as a seemingly constitutive complex that undergoes self-assembly into fibres of many micrometres in length (*29*), which likely provide the long-range structural supports that stabilise continuous growth of the SYCP1 lattice along the entire chromosome axis (*22*).

Owing to the essential roles of meiotic recombination, synapsis and chromosome dynamics in mammalian meiosis (*15, 30–34*), their defects are associated with human infertility, recurrent miscarriage and aneuploidies (*35, 36*). As genetic causes of infertility, they typically fall within the category of idiopathic cases, having no readily diagnosable and clinically resolvable cause. Within the 10-15% of couples who suffer from infertility, approximately 25% are idiopathic and of likely genetic origin, comprising 50-80% of cases of non-obstructive azoospermia (NOA) and premature ovarian failure (POF) (*36, 37*). Whilst individual infertility mutations are inherently unlikely to become widespread in a population, they can be found within families, especially when consanguineous (*38*), and provide crucial insights into their common targets and the molecular mechanisms that they disrupt.

Within the SC, familial infertility mutations have been identified for SYCP3 and SYCE1 (*36*). All identified SYCP3 mutations are autosomal dominant and alter or delete its structural core’s C-terminus that mediates filamentous assembly, so likely sequester wild-type molecules into inactive complexes (*24, 36*). In contrast, the three identified SYCE1 mutations are autosomal recessive and were found in two familial cases of NOA and one of POF (*36*). The two NOA cases are splice-site mutations, c.197-2A>G and c.375-2A>G, which are predicted to result in a truncated product of amino-acids 1-65 and an internal deletion of amino-acids 126-155, respectively (*39, 40*). These remove or delete part of human SYCE1’s structural core that is encoded by amino-acids 25-179, so can be explained by disruption of its dimeric structure (Figure 1b) (*28, 36*). The POF mutation, c.613C>T, generates a premature stop codon (p.Gln241*) to give a truncated product of amino-acids 1-240, relative to the canonical 351 amino-acids isoform (Figure 1b) (*41*). However, as this truncation lies outside SYCE1’s structural core, the molecular mechanism that is disrupted, and thereby responsible for infertility, remains unknown.

Here, we combine mouse genetics, cellular and biochemical studies to reveal a multivalent interaction mode between SYCE1 and SIX6OS1 that is disrupted by infertility-associated mutations of SYCE1. We find that the SIX6OS1 N-terminus binds and disrupts the core dimeric structure of SYCE1 (amino-acids 25-179) to form a 1:1 complex as the first interface, and its downstream sequence binds to SYCE1 amino-acids 177-305 as the second interface. SYCE1’s infertility-associated mutations c.375-2A>G (NOA) and c.613C>T (POF) specifically disrupt the first and second interfaces, respectively. Mice harbouring the SYCE1 POF mutation and a targeted deletion within SIX6OS1 (which disrupts the first interface) are infertile, with failure of SC assembly. We conclude that both SYCE1-SIX6OS1 binding interfaces are essential for SC assembly and meiotic division, thus explaining how human infertility results from the differential targeting of binding interfaces by SYCE1’s reported clinical mutations.

## Results

### SYCE1 premature ovarian failure mutation c.613C>T retains its core dimeric structure

The SYCE1 premature ovarian failure mutation c.613C>T encodes a premature stop codon (p.Gln241*) that is predicted to generate a truncated protein product of amino-acids 1-240, relative to SYCE1’s canonical 351 amino-acids isoform (Figure 1b) (*41*). We previously demonstrated that an N-terminal structural core encoded by amino-acids 25-179 (SYCE1core) forms an α-helical anti-parallel coiled-coil structure that mediates head-of-head dimerization of SYCE1 (*28*). As this core region is retained (Figure 1b), we predicted that SYCE1’s anti-parallel dimeric structure would be maintained within the 1-240 truncated product of the POF mutation (SYCE1POF). To test this, we purified recombinant SYCE1POF, generating purified material that contained approximately equal quantities of the full protein and a degradation product of apparent size consistent with C-terminal degradation to its structural core (Figure 1c). Circular dichroism (CD) spectroscopy confirmed that SYCE1POF contains a proportion of α-helical structure consistent with retention of the 25-179 core structure (Figure 1d), and SYCE1POF and SYCE1core demonstrated identical melting temperatures (*T_m_*) of 39°C (Figure 1e). Further, analysis by size-exclusion chromatography multi-angle light-scattering (SEC-MALS) confirmed that the full and degraded proteins are homodimers of 48 kDa and 39 kDa, respectively (Figure 1f). We conclude that SYCE1POF retains the dimeric structure imposed by its core 25-179 region, so its SC and meiotic defects must result from additional structural or functional roles of its deleted C-terminus.

### The SYCE1 POF mutation leads to failure of SC assembly and infertility in mice

Having established its retention of core dimeric structure, we next sought to determine the structural and functional consequence of the SYCE1 POF mutation on the SC and meiotic division *in vivo*. We thus generated mice harbouring mutations of *Syce1* alleles to introduce stop codons at amino-acid position 243, equivalent to the human p.Gln241* mutation (Supplementary Figure 1). Whilst heterozygotes (designated *Syce1^POF/WT^*) were fertile, both male and female homozygotes (designated *Syce1^POF/POF^*) were infertile, replicating the autosomal recessive pattern of the POF mutation in humans (*41*). In male mutant mice, we observed reduced testis size (Figure 2a) and a pachytene-like arrest similar to that observed in the SYCE1 knockout (*16*). There was defective SC assembly, with reduced staining for SYCP1 (Figure 2b) and SYCE3 (Figure 2c) and no staining for SYCE1 (Figure 2d), SIX6OS1 (Figure 2e) and SYCE2-TEX12 (Figures 2f and 2g). We next studied the kinetics of DSB repair. Meiotic DSBs are generated by the nuclease SPO11 and are then resected to form ssDNA ends that invade into the homologous chromosome by the recombinases RAD51 and DMC1 (*42*). DSBs are labelled by the presence of phosphorylated H2AX (γ-H2AX) (*43*). The distribution of γ-H2AX in mutant spermatocytes was similar to that found in WT cells at early prophase I but show increased staining at pachytene-like arrest (Supplementary Figure 2a). The distributions of RAD51 and DMC1 were detected on aligned LEs (Supplementary Figures 2b and 2c) but in absence of mismatch repair protein MLH1 (marker of crossing overs) (Supplementary Figure 2d). Together, these data indicate proper generation of DSBs but with failure of their repair and CO formation in *Syce1^POF/POF^*. In female mutant mice, we observed no follicles in adult ovaries (Figure 3a), and embryonic oocytes demonstrated pachytene arrest with mostly unaligned chromosome axes, recapitulating the human POF syndrome. Analysis of the SC revealed similar defects, with reduction in SYCP1 (Figure 3b) and SYCE3 (Figure 3c) staining (though to a lesser extent than males), and absence of SYCE1 (Figure 3d), SIX6OS1 (Figure 3e) and SYCE2-TEX12 (Figures 3f and 3g). The distribution of γ-H2AX, RAD51 and DMC1 labelling in pachynema-like mutant oocytes was also increased and lacked MLH1 foci (Supplementary Figures 3a-d). Thus, the SYCE1 POF mutation leads to male and female infertility with phenotypes of failed DSB repair, synapsis and finally SC assembly, similar to those previously observed upon disruption of structural components of the SC central element (*10, 11, 16-18*).

**Figure 2.**
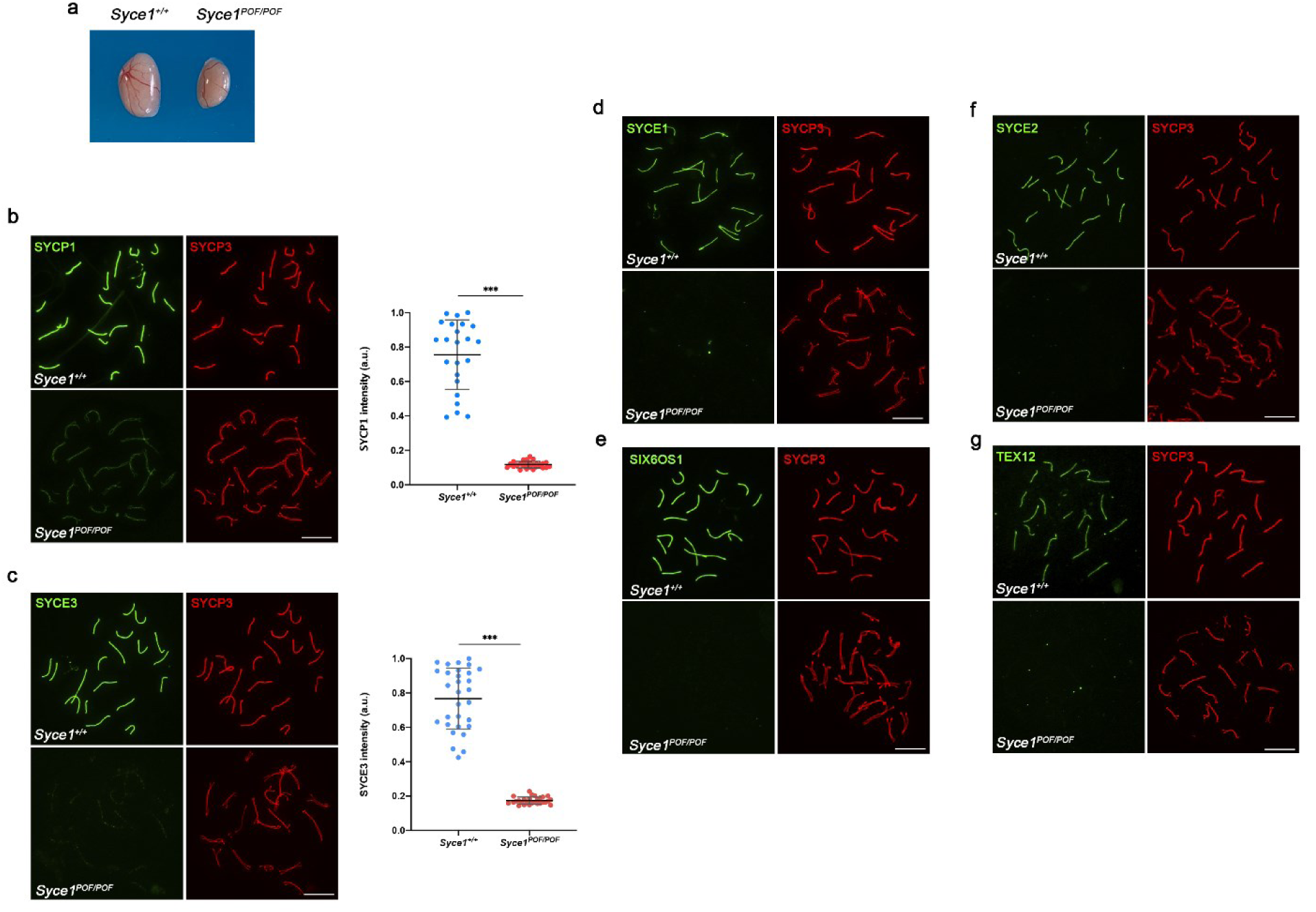
Characterization of *Syce1^POF/POF^* mutant mice, whose spermatocytes are not able to synapse. **(a)** Testis from WT and *Syce1^POF/POF^* mice showing reduced testis size. **(b)** Double immunolabeling of WT pachytene and *Syce1^POF/POF^* pachytene-like spermatocytes with SYCP3 (red) and SYCP1 (green). In *Syce1^POF/POF^* spermatocytes AEs fail to synapse and show a weak staining of SYCP1 along the AEs. **(c-g)** Double immunolabeling of spermatocyte spreads with SYCP3 (red) and the CE proteins (green). *Syce1^POF/POF^* pachytene-like spermatocytes show a highly reduced signal of SYCE3 **(c)** and the absence of all **(d)** SYCE1, **(e)** SIX6OS1, **(f)** SYCE2 and **(g)** TEX12 from the AEs. Scale bars represent 10 μm. Plot right to the panels **(b-c)** represent the quantification of the fluorescence intensity levels in *Syce1^POF/POF^* pachytene-like spermatocytes compared to WT pachytenes. Welch’s *t*-test analysis: * p<0.01, **p<0.001, ***p<0.0001.

**Figure 3.**
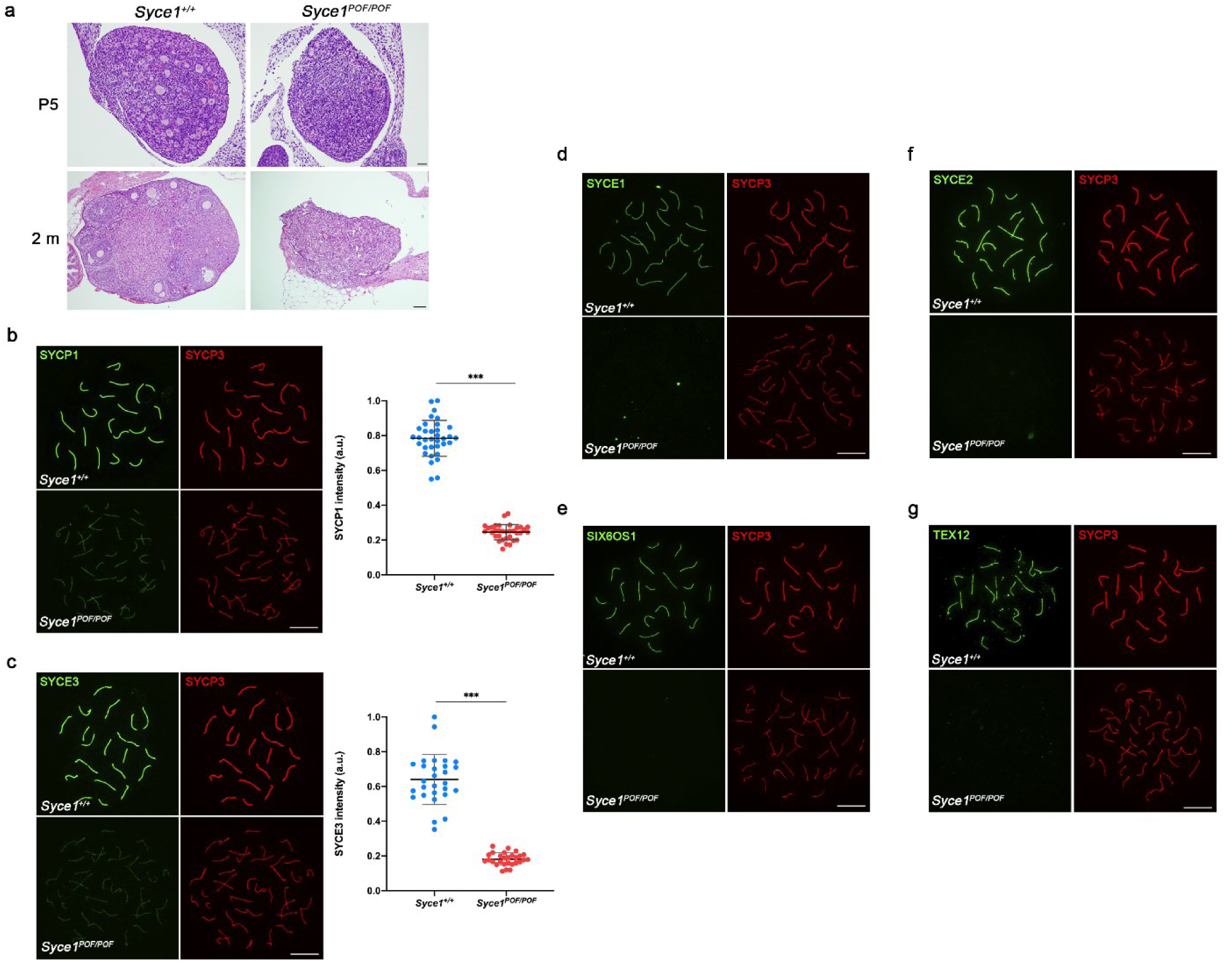
*Syce1^POF/POF^* oocytes fail to synapse. **(a)** Hematoxylin-eosin staining of sections of ovaries from WT and *Syce1^POF/POF^* female mice showing depletion of follicles at 5 days (P5) and 2 months of age (2m). Scale bars represent 20 μm in P5 and 50 μm at 2m ovaries. **(b)** Double immunolabeling of oocyte spreads from WT and *Syce1^POF/POF^* mice with SYCP3 (red) and SYCP1 (green). *Syce1^POF/POF^* oocytes get arrested in a pachytene-like stage where AEs are not able to synapse and show a marked unalignment, with reduced levels of SYCP1. **(c-g)** Double immunolabeling of oocyte spreads with SYCP3 (red) and the CE proteins (green). *Syce1^POF/POF^* pachytene-like oocytes show reduced signal of SYCE3 **(c)** and a complete absence of all **(d)** SYCE1, **(e)** SIX6OS1, **(f)** SYCE2 and **(g)** TEX12 from the AEs. Scale bars represent 10 μm. Plots right to the panels **(b-c)** represent the quantification of the fluorescence intensity levels in *Syce1^POF/POF^* pachytene-like oocytes compared to WT pachytenes. Welch’s *t*-test analysis: * p<0.01, **p<0.001, ***p<0.0001.

### SYCE1POF retains SIX6OS1-binding but lacks SYCE3-binding in heterologous systems

As the *Syce1^POF/POF^* mouse strain indicated a clear structural defect in the SC, we wondered whether the POF mutation may disrupt the known interaction between SYCE1 and fellow SC central element components SIX6OS1 and SYCE3 (*11*). The expression of SYCE1 and SIX6OS1 in COS7 cells produced cytoplasmic signals that became co-localised in foci upon co-expression (Figure 4a), in keeping with our previous findings (*11*). SYCE1POF formed similar or slightly reduced numbers of foci that equally co-localized with SIX6OS1, indicating a retention of SIX6OS1-binding (Figure 4a). We further demonstrated a similar co-immunoprecipitation of SIX6OS1 by wild-type SYCE1 and SYCE1POF upon co-expression in HEK293 cells (Figure 4b). Thus, the SYCE1-SIX6OS1 interaction is retained in the SYCE1 POF mutation. Could other disrupted functions contribute to the effect of the POF mutation? The only other known SYCE1 interactor is SYCE3, which undergoes low-affinity binding (Crichton *et al*, manuscript in preparation). In contrast with the wild-type protein, the expression of SYCE1POF (cytoplasmic foci) in COS7 cells failed to recruit SYCE3 (preferentially nuclear) to their cytoplasmic foci (loss of co-localization, Figure 4c). Similarly, SYCE1POF failed to co-immunoprecipitate SYCE3 upon co-expression in HEK293 cells (Figure 4d). Thus, whilst the high-affinity SYCE1-SIX6OS1 complex is retained, the low-affinity SYCE1-SYCE3 complex is largely abolished in the SYCE1 POF mutation.

**Figure 4.**
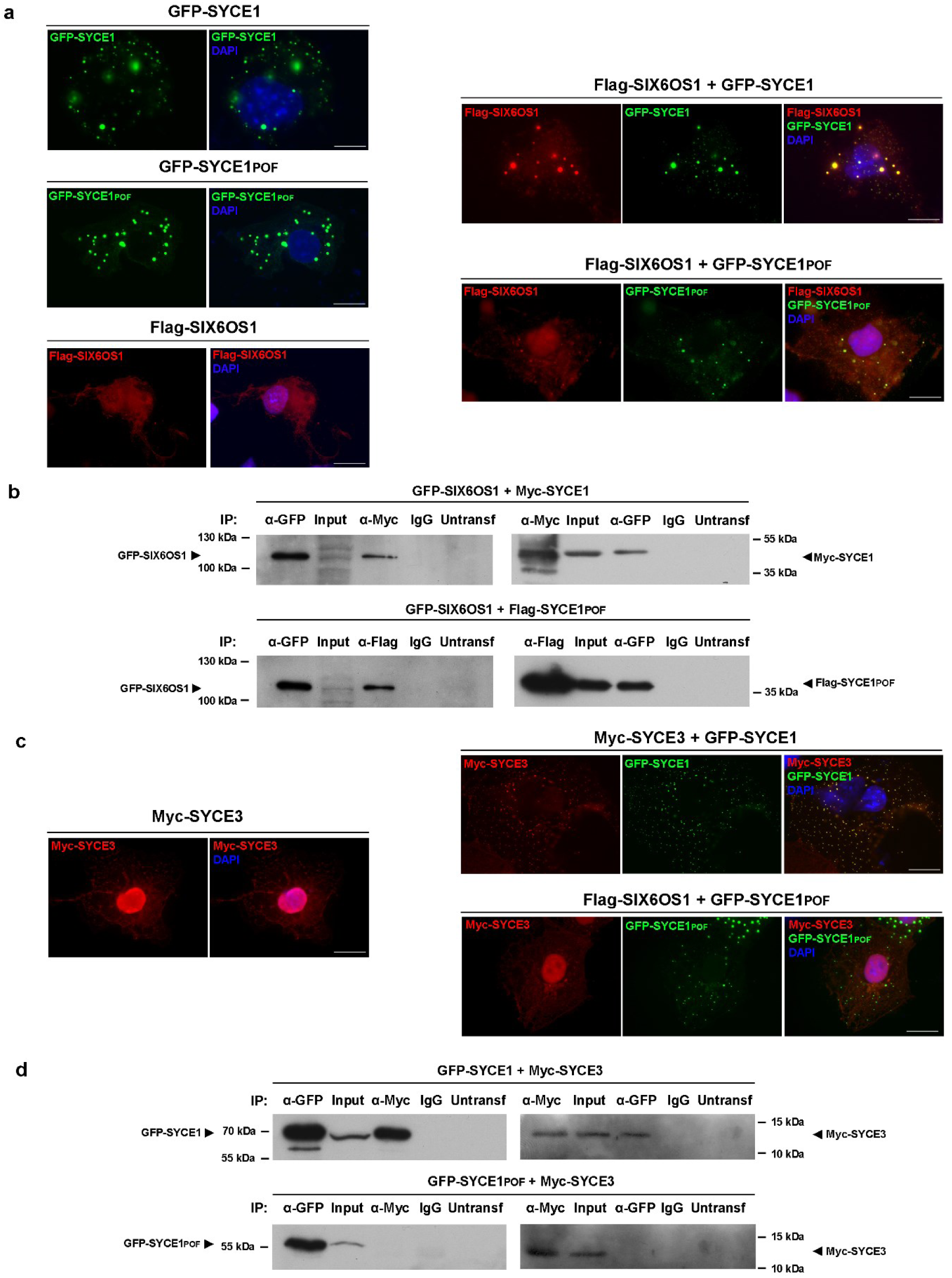
SYCE1POF retains SIX6OS1-binding but fails to retain the SYCE3-interaction in heterologous systems. (**a**) COS7 cells were transfected or co-transfected with *Syce1*, *Syce1POF* and *Six6os1*, either alone (left panel) or in different combinations (right panel), as indicated. SIX6OS1 was found to co-localize with both SYCE1 and SYCE1POF in the cytoplasmatic punctate pattern of SYCE1 upon co-expression. Scale bars represent 20 µm. (**b**) HEK 293T cells were co-transfected with the indicated expression vectors. Protein complexes were immunoprecipitated overnight with either an anti-Flag, anti-EGFP or anti-Myc antibody, and were analysed by immunoblotting with the indicated antibody. GFP-SIX6OS1 co-immunoprecipitates with both Myc-SYCE1 and Flag-SYCE1POF (as well as in the reciprocal IP), suggesting that the POF mutation of SYCE1 alone is not enough for the disturbance of the interaction. (**c**) COS7 cells were transfected with *Syce3* either alone (left panel) or in combination with *Syce1* and *Syce1POF* (right panel), as indicated. SYCE1 showed to co-localize with SYCE3 in his own cytoplasmatic punctate pattern. The co-localization is heavily hampered with SYCE1POF. Scale bars represent 20 µm. (**d**) HEK 293T cells were co-transfected with the indicated expression vectors. Protein complexes were immunoprecipitated overnight with either an anti-Myc or anti-EGFP antibody and were analysed by immunoblotting with the indicated antibody. SYCE1 co-immunoprecipitates SYCE3 (as well as in the reciprocal IP), but the interaction is eradicated in the SYCE1 POF condition, suggesting that the C-terminal region of SYCE1 is required for the interaction with SYCE3.

### SYCE1core undergoes conformational change to form a 1:1 complex with SIX6OS1

What is the molecular basis of SIX6OS1-binding by SYCE1? As this is retained in SYCE1POF, we reasoned that SIX6OS1-binding must be mediated by SYCE1’s structural core. We screened SYCE1core against a library of SIX6OS1 constructs through bacterial co-expression and identified a robust interaction with amino-acids 1-67 of SIX6OS1, herein referred to as SIX6OS1N (Figures 1b and 5a). We were able to purify the SYCE1core-SIX6OS1N complex by reciprocal affinity chromatography, ion exchange and size-exclusion chromatography (Figure 5b) and identified no biochemical conditions in which it was disrupted, indicating that it is a high-affinity interaction. We were further able to purify similar complexes for SYCE1POF (with the same degradation product as upon isolated expression) and full-length SYCE1 (Figure 1c), confirming that SIX6OS1-binding is retained by all constructs containing the 25-179 core. CD analysis revealed similar α-helical content for SYCE1-SIX6OS1N complexes as for their isolated SYCE1 proteins (Figure 1d). CD thermal denaturation revealed slightly increased cooperativity of unfolding and melting temperatures for SYCE1-SIX6OS1N complexes relative to their isolated SYCE1 proteins (increasing from 39°C to 43°C, 39°C to 41°C and 38°C to 40°C for SYCE1core, SYCE1POF and full-length, respectively; Figures 1e and 5c). Remarkably, SEC-MALS analysis revealed that all three SYCE1-SIX6OS1N complexes are 1:1, with molecular weights of 27 kDa, 37 kDa and 46 kDa, respectively (Figure 5d and Supplementary Figure 4a). Thus, the SYCE1core undergoes conformation change from an anti-parallel homodimer to a 1:1 complex upon binding to SIX6OS1N (Figure 5e).

**Figure 5.**
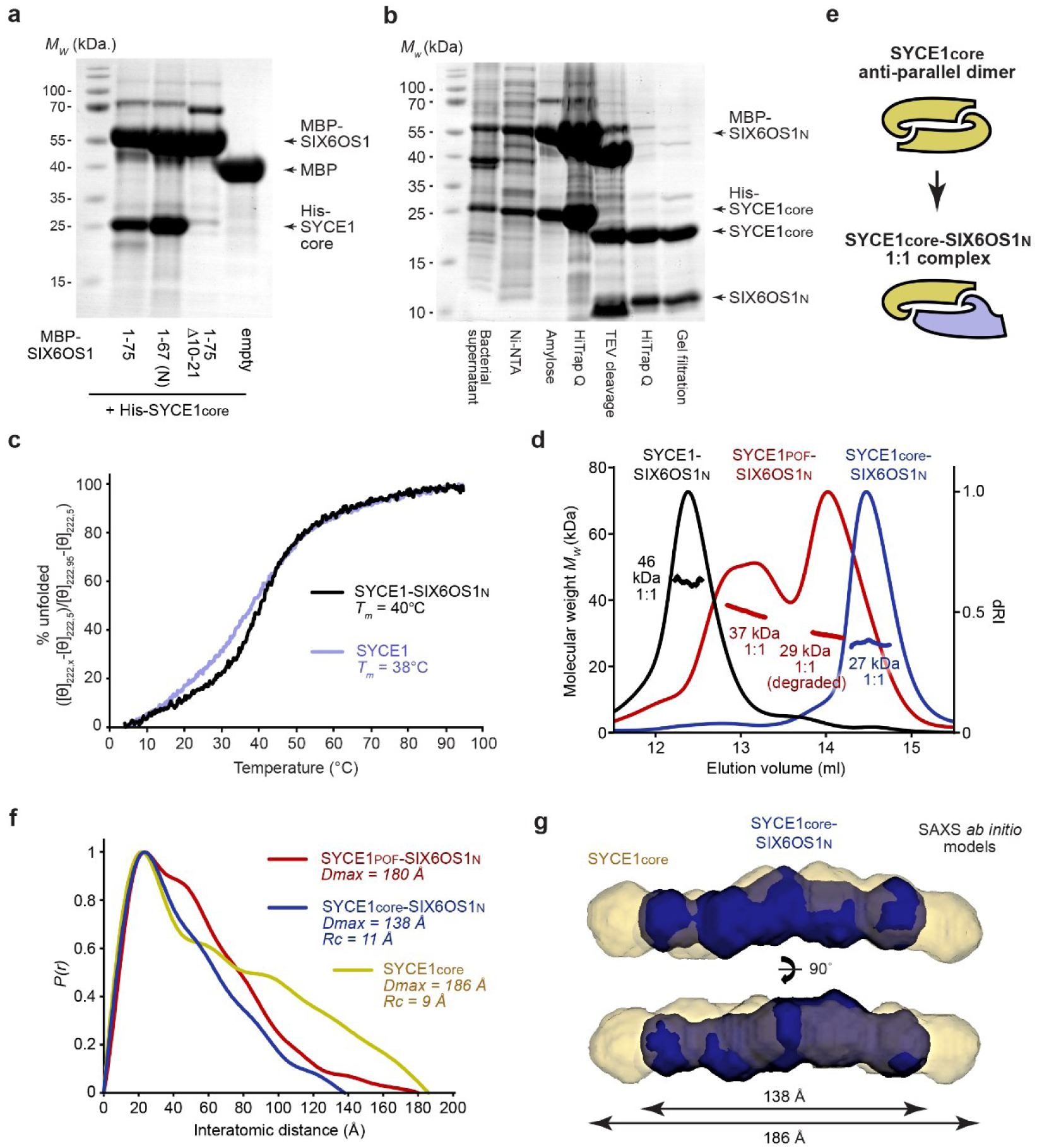
SYCE1core undergoes conformational change to form a 1:1 complex with SIX6OS1N. (**a**) Amylose pull-down following co-expression of MBP-SIX6OS1 1-75, 1-67, 1-75 Δ10-21 and free MBP with His-SYCE1core. (**b**) SDS-PAGE analysis of the co-expression and co-purification of the SYCE1core-SIX6OS1N complex; recombinant proteins were purified through amylose and anion exchange chromatography, followed by TEV cleavage to remove N-terminal His- and MBP-tags, with subsequent anion exchange and size exclusion chromatography. (**c**) CD thermal denaturation recording the CD helical signature at 222 nm between 5°C and 95°C, as % unfolded; estimated melting temperatures (*Tm*) are indicated. (**d**) SEC-MALS analysis. SYCE1core-SIX6OS1N (blue), SYCE1POF-SIX6OS1N (red) and full-length SYCE1-SIX6OS1N (black) are 1:1 complexes of 27 kDa, 37 kDa (29 kDa for the degradation product complex) and 46 kDa, respectively (theoretical 1:1 – 27 kDa, 36 kDa and 48 kDa). (**e**) Schematic of the conformational change of the SYCE1core anti-parallel dimer (yellow) into a 1:1 SYCE1core-SIX6OS1N complex (yellow-blue). (**f,g**) Size-exclusion chromatography small-angle X-ray scattering (SEC-SAXS) analysis. (**f**) SEC-SAXS *P(r)* interatomic distance distributions of SYCE1core-SIX6OS1N (blue), SYCE1POF-SIX6OS1N (red) and SYCE1core (yellow), revealing maximum dimensions (*Dmax*) of 138 Å, 180 Å and 186 Å, respectively. Their cross-sectional radii (*Rc*) are indicated (Supplementary Figure 4d). (**g**) SAXS *ab initio* models of SYCE1core-SIX6OS1N (blue) and SYCE1core (yellow); averaged models were generated from 20 independent DAMMIF runs with a NSD values of 0.730 (± 0.0546) and 0.726 (± 0.058) and reference model χ^2^ values of 0.651 and 1.067. Data for SYCE1core and full-length SYCE1 are reproduced from Dunne and Davies (*28*).

We analysed the conformation of the SYCE1core-SIX6OS1N complex by size-exclusion chromatography small-angle X-ray scattering (SEC-SAXS; Supplementary Figures 4b and 4c). The SAXS real space pair-distance *P(r)* distribution (the distribution of interatomic distances within a protein structure) demonstrates positive skew, indicating that SYCE1core-SIX6OS1N retains the rod-like structure of SYCE1core, but with a reduction in its molecular length from 186 Å to 138 Å (Figure 1f). Further, its cross-sectional radius is slightly increased from 9 Å to 11 Å (Supplementary Figure 4d), suggesting an increase from a two- to four-helical coiled-coil. These geometric changes are consistent with the SYCE1core-SIX6OS1N 1:1 complex forming a shorter but wider coiled-coil than the isolated SYCE1core dimer, as indicated by their SAXS *ab initio* models (Figure 5g). Further, the SAXS *P(r)* distribution of SYCE1POF indicates a similar elongated structure but with an increased tail to a maximum dimension of 180 Å (Figure 5f), consistent with it containing the same SYCE1core-SIX6OS1N structure with an extended and potentially unstructured C-terminus to amino-acid 240. We conclude that SYCE1core mediates a direct interaction with SIX6OS1N that imposes a conformational change to a 1:1 complex that adopts a shorter and wider coiled-coil conformation than the isolated SYCE1core anti-parallel homodimer.

### SYCE1POF disrupts a second SYCE1-SIX6OS1 binding interface

Does the SYCE1core-SIX6OS1N complex represent the sole means by which SYCE1 interacts with SIX6OS1? We were unable to obtain soluble biochemical complexes containing SIX6OS1 sequences beyond its N-terminus, so utilised yeast two-hybrid (Y2H) to test SYCE1-binding by full-length SIX6OS1. Having confirmed direct binding of SYCE1core to full-length SIX6OS1, we used C-terminal truncation to dissect its minimal binding site to amino-acids 1-75, in keeping with our biochemical findings, and identified an additional interaction between SYCE1 177-305 and full-length SIX6OS1 (Figure 6a).

**Figure 6.**
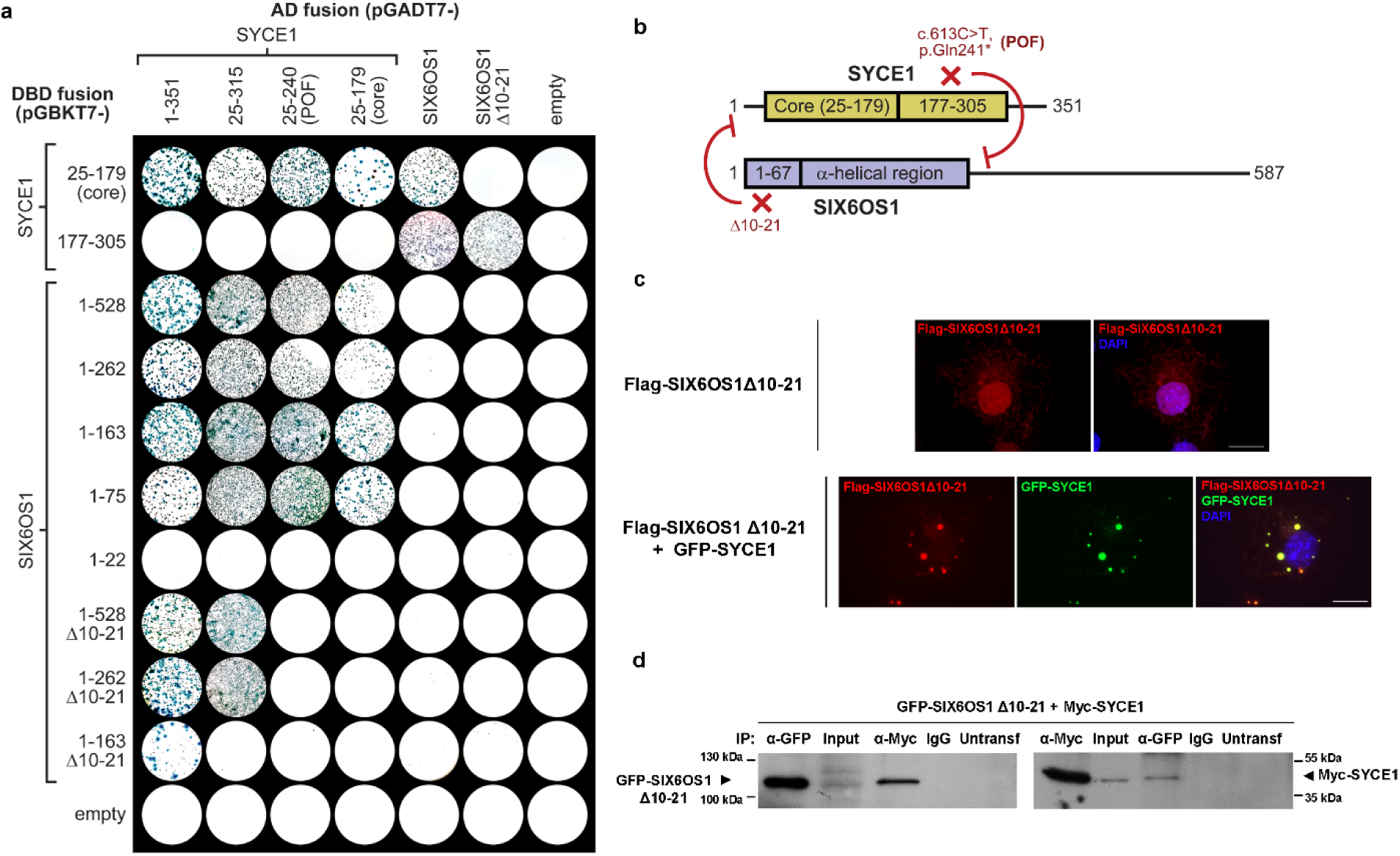
SYCE1 undergoes multivalent interaction with SIX6OS1 in yeast but SIX6OS1 Δ10-21 retains SYCE1-binding in heterologous systems. (**a**) Yeast two hybrid analysis of interactions between SYCE1 and SIX6OS1, using the constructs indicated. Y187 and Y2HGold yeast strains harbouring pGBKT7 and pGADT7 plasmids were mated and plated onto SD/-Ade/-His/-Leu/-Trp/X-α-Gal. Positive reactions depend on activation of reporter genes *ADE1*, *HIS3* and *MEL1*. These data are representative of three repeats. (**b**) Schematic of the SYCE1-SIX6OS1 interaction highlighting that the SYCE1 POF mutation blocks the second binding interface between SYCE1 177-305 and SIX6OS1 downstream sequence within region 1-262, whereas the SIX6OS1 Δ10-21 deletion blocks the first binding interface between SYCE1core (25-179) and SIX6OS1N (1-67). (**c**) COS7 cells were transfected with *Six6os1 Δ10-21* either alone (upper panel) or in combination with *Syce1* (lower panel). When transfected alone, SIX6OS1 Δ10-21 shows a change in its cytological localization from the typical wild type cytoplasmatic pattern to nuclear localization in addition to some remaining cytoplasmatic signal. Furthermore, SIX6OS1 Δ10-21 retained its ability to form co-localized foci with SYCE1 when co-transfected. Scale bars represent 20 µm. (**d**) HEK 293T cells were co-transfected with the indicated expression vectors. Protein complexes were immunoprecipitated overnight with either an anti-Myc or anti-EGFP antibody and were analysed by immunoblotting with the indicated antibody. SIX6OS1 Δ10-21 retains its ability to co-immunoprecipitate SYCE1 (as well as in the reciprocal IP), suggesting that the second SYCE1-SIX6OS1 binding interface is preserved in SIX6OS1 Δ10-21.

In order to establish whether SYCE1core and 177-305 bind to the same or distinct sites within SIX6OS1, we established an internal deletion of SIX6OS1 amino-acids 10-21 (Δ10-21) that blocks formation of the SYCE1core-SIX6OS1N biochemical complex (Figure 6a). Remarkably, whilst Δ10-21 completely abrogated the Y2H interaction of full-length SIX6OS1 with SYCE1core (25-179), it retained a robust interaction with SYCE1 177-305, suggesting distinct SIX6OS1-binding sites. Further, Δ10-21 blocked the ability of SIX6OS1 1-262 to interact with SYCE1core and SYCE1POF (amino-acids 25-240) whilst retaining its binding to full-length and 25-315 SYCE1 (Figure 6a). Thus, SYCE1 undergoes multivalent interactions with SIX6OS1, with the first binding interface mediated by SYCE1core and SIX6OS1N (1-67), and the second interface mediated by SYCE1 177-305 and downstream sequence within SIX6OS1 1-262. Further, the first and second binding interfaces are specifically disrupted by SIX6OS1 deletion Δ10-21 and the SYCE1 POF mutation, respectively, and in both cases an SYCE1-SIX6OS1 complex is retained through the unaffected alternative site (Figure 6b).

### SIX6OS1 Δ10-21 retains SYCE1-binding in heterologous systems

Our biochemical and Y2H analyses concluded that SIX6OS1 Δ10-21 would disrupt the first SYCE1-SIX6OS1 binding interface whilst retaining complex formation through the second interface. In support of this, we found that SIX6OS1 Δ10-21 retained its ability to form intense co-localised foci with SYCE1 upon co-expression in COS7 cells (Figure 6c), similar to our previous observations for the SYCE1 POF mutation. Similarly, SIX6OS1 Δ10-21 retained its ability to co-immunoprecipitate SYCE1 upon co-expression in HEK293 cells (Figure 6d). Thus, localisation and co-immunoprecipitation data from heterologous systems support our Y2H findings that the second SYCE1-SIX6OS1 binding interface is retained in SIX6OS1 Δ10-21, mirroring the retention of only the second binding interface that is predicted for the 126-155 deletion of the SYCE1 c.375-2A>G NOA mutation (*40*).

### SIX6OS1 Δ10-21 leads to failure of SC assembly and murine infertility

Having established that the severe phenotype of the SYCE1 POF mutation likely results from the sole disruption of the second SYCE1-SIX6OS1 binding interface, we wondered whether a similar phenotype would result from the sole disruption of the first interface. To test this, we generated mice harbouring mutations of *Six6os1* alleles encoding internal in-frame deletions of amino-acids 10-21 (equivalent numbering to the human protein) (Supplementary Figures 5a and 5b). Whilst heterozygotes (designated *Six6os1^Δ10-21/WT^*) were fertile, both male and female homozygotes (designated *Six6os1*^Δ*10-21/*Δ*10-21*^) were infertile, similar to the SYCE1 POF mutation. In males, we observed reduced testis size (Figure 7a) and a pachytene-like arrest similar to that observed in the *Six6os1* and *Syce1* knockouts (*11, 16*). The mutant spermatocytes were defective in synapsis and SC assembly, with reduced staining for SC proteins SYCP1 (Figure 7b), SYCE3 (Figure 7c), and no staining for SYCE2-TEX12 (Figures 7f and 7g). In contrast with their complete absence in the SYCE1 POF mutation, we observed some residual staining for SYCE1 (Figure 7d) and SIX6OS1 (Figure 7e). We detected γH2AX (Supplementary Figure 6a) and DMC1/RAD51 foci (Supplementary Figures 6b and 6c) on aligned axial elements but no MLH1 foci (Supplementary Figure 6d), indicating the proper induction of DSBs with their failed repair and absence of COs. Thus, SIX6OS1 Δ10-21 leads to infertility with a phenotype of failed DSB repair and SC assembly, similar to the SYCE1 POF mutation and those reported for disruption of structural components of the CE (*10, 11, 16-18*).

**Figure 7.**
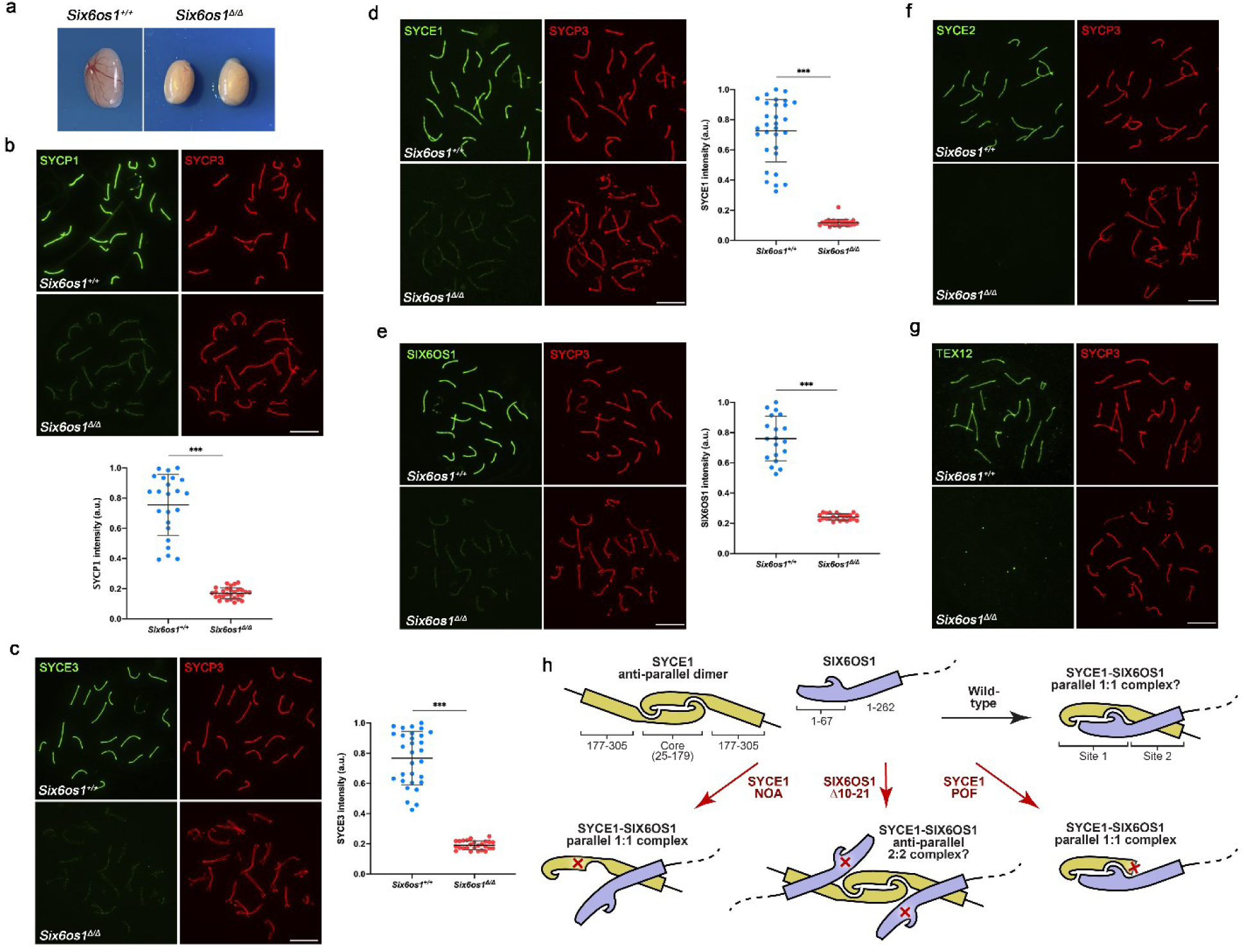
Synapsis between homologues is disrupted in *Six6os1*^Δ*10-21/*Δ^*^10-21^* spermatocytes. **(a)** Genetic deletion of amino-acids 10-21 of SIX6OS1 leads to a reduction of the testis size comparted to the WT. **(b)** Double immunolabeling of WT pachytene and *Six6os1*^Δ*/*Δ^ pachytene-like spermatocytes with SYCP3 (red) and SYCP1 (green). AEs fail to synapse in *Six6os1*^Δ*/*Δ^ spermatocytes despite the partial alignment with reduced loading of SYCP1 along the AEs. **(c-g)** Double immunolabeling of spermatocyte spreads with SYCP3 (red) and all the CE components (green). *Six6os1*^Δ*/*Δ^ pachytene-like spermatocytes show reduced signal of **(c)** SYCE3, **(d)** SYCE1 and **(e)** SIX6OS1 coupled with the absence of **(f)** SYCE2 and **(g)** TEX12 from the AEs. Scale bars represent 10 μm. Plots represent the quantification of the fluorescence intensity levels in *Six6os1*^Δ*/*Δ^ pachytene-like spermatocytes compared to WT pachytenes **(b-e)**. Welch’s *t*-test analysis: * p<0.01, **p<0.001, ***p<0.0001. (**h**) Schematic of how the SYCE1 anti-parallel dimer (yellow) undergoes conformational change upon interaction with SIX6OS1 (blue) to form a possible 1:1 complex through consecutive binding interfaces mediated by SYCE1core-SIX6OS1N (site 1) and SYCE1 177-305 and downstream sequence within SIX6OS1 1-262 (site 2). The consequence of SYCE1 mutations associated with POF (c.613C>T) and NOA (c.375-2A>G) and SIX6OS1 Δ10-21 on the integrity of the two binding interfaces, and the predicted stoichiometry and conformation of resultant SYCE1-SIX6OS1 complexes, is illustrated.

Thus, we conclude that both first and second SYCE1-SIX6OS1 binding interfaces are essential for SC assembly and meiotic progression. Further, these findings explain how the sole disruption of individual SYCE1-SIX6OS1 binding interfaces by SYCE1 NOA (c.375-2A>G) and POF (c.613C>T) mutations result in the reported familial cases of human infertility.

## Discussion

The structural and functional integrity of the SC is contingent on the structure and assembly of is constituent protein components. Here, we report that SC assembly depends on multivalent interactions between central element components SYCE1 and SIX6OS1 that are disrupted by infertility-associated mutations of SYCE1. The first binding interface is formed by the structural core of SYCE1 (SYCE1core; amino-acids 25-179), which undergoes conformational change from an anti-parallel homodimer to a 1:1 complex upon interaction with SIX6OS1’s N-terminus (SIX6OS1N; amino-acids 1-67). The second binding interface is formed by downstream sequence within SIX6OS1 1-262 interacting directly with SYCE1 177-305. Through the generation of mice harbouring an internal deletion of SIX6OS1’s N-terminus (Δ10-21) and the SYCE1 POF mutation (murine p.Gln243*), which specifically block the first and second binding interfaces, respectively, we find that integrity of both SYCE1-SIX6OS1 binding interfaces is essential for SC assembly and meiotic progression *in vivo*.

What is the structure of the SYCE1-SIX6OS1 complex? SEC-SAXS analysis revealed that the SYCE1core-SIX6OS1N 1:1 complex formed by the first binding interface has length and cross-sectional radius of 138 Å and 11 Å, in comparison with 186 Å and 9 Å for the SYCE1core dimer. We previously reported a model for SYCE1core in which amino-acids 52-179 form an anti-parallel dimeric coiled-coil containing a midline ‘kink’, with α-helices of amino-acids 25-50 packing against this structural core (*28*). A maximum dimension of 138 Å for SYCE1core-SIX6OS1N suggests a coiled-coil length of approximately 92 amino-acids, given a helical rise of 1.5 Å per amino-acid (*44*). This could be explained by the 52-179 region forming a helix-turn-helix structure through exaggeration of the ‘kink’ to a full turn, which may combine with the α-helix formed by amino-acids 25-50 and an α-helix from SIX6OS1N to form a four-helical coiled-coil, consistent with its 11 Å cross-sectional radius. The second binding interface between SYCE1 177-305 and downstream sequence within SIX6OS1 1-262 suggests that SYCE1core-SIX6OS1N likely adopts a parallel configuration to form a single SYCE1-SIX6OS1 1:1 complex of consecutive first and second binding interfaces (Figure 7h).

Our analysis of the SYCE1-SIX6OS1 complex reveals how the three reported clinical mutations of SYCE1 differentially affect its interaction with SIX6OS1. The SYCE1 NOA mutation c.197-2A>G is predicted to result in a truncated product of amino-acids 1-65 (*39*), which would disrupt both binding sites so likely abrogates SYCE1-SIX6OS1 complex formation and thus works as a null mutation. The SYCE1 NOA mutation c.375-2A>G is predicted to result in internal deletion of amino-acids 126-155 (*40*), which would disrupt the first binding interface whilst retaining the second binding interface, so is likely to result in a conformationally-altered 1:1 complex (Figure 7h). In contrast, whilst Δ10-21 SIX6OS1 similarly disrupts the first binding interface and retains the second binding interface, the SYCE1core remains unaffected so is predicted to enable formation of a head-to-head 2:2 complex (Figure 7h). The SYCE1 POF mutation c.613C>T generates a premature stop codon (p.Gln241*) that gives a truncated product of amino-acids 1-240 (*41*), which we have demonstrated disrupts the second binding interface whilst retaining the first binding interface (Figure 7h). Thus, the latter two infertility-associated mutations of SYCE1 specifically disrupt one SYCE1-SIX6OS1 interface whilst retaining the other, which combine with our mouse genetic studies to confirm that both interfaces are essential for the structural assembly of the SC and its function in meiosis.

What are the structural roles of SYCE1 and SYCE1-SIX6OS1 within the SC? Our analyses of *Syce1^POF/POF^* and *Six6os1^Δ10-21/ Δ10-21^* mouse strains revealed similar phenotypes with retention of some SYCP1 and SYCE3 recruitment to chromosome axes, with absence or substantial reduction of SYCE1 and SIX6OS1, and lack of recruitment of SYCE2-TEX12. This pattern suggests a hierarchical model of SC assembly in which SYCE1 and SYCE1-SIX6OS1 lie downstream of SYCP1 and SYCE3, and upstream of SYCE2-TEX12. This is consistent with knockout data (*10, 11, 15-18*) and with our unpublished findings that SYCP1 forms a high-affinity complex with SYCE3, which provides the sole link with SYCP1 and recruits other CE proteins via low-affinity interactions with SYCE1 and SYCE2-TEX12 (Crichton *et al*, manuscript in preparation). The disruption of SYCE3-binding by the POF mutation suggests that its SYCE1-SIX6OS1 complex would be defective for SC recruitment, whereas the SYCE1-SYCE3 interaction, and hence SC recruitment, should be retained for the SYCE1-SIX6OS1 complex of the SIX6OS1 Δ10-21 internal deletion. This explains the greater severity of the CE loading defect in *Syce1^POF/POF^* than *Six6os1^Δ10-21/ Δ10-21^*, in which SYCE1 and SIX6OS1 staining were substantially reduced in the latter but completely absent in the former. Thus, we conclude that the first and second SYCE1-SIX6OS1 interfaces are essential for initiation of SC central element formation, and likely function by stabilising a local three-dimensional SC structure which mediates recruitment and self-assembly of SYCE2-TEX12 into fibres that mediate SC elongation along the chromosome axis. Further, the SYCE1 POF mutation is likely worsened by its additional disruption of SYCE3-binding that removes the residual SYCE1-SIX6OS1 SC recruitment observed for the SIX6OS1 Δ10-21 internal deletion.

The existence of SYCE1core as an isolated anti-parallel homodimer and in a 1:1 complex with SIX6OS1N raises the question of which is the biologically-relevant conformation? It is important to highlight that the CD melting temperatures of SYCE1-SIX6OS1N complexes and isolated SYCE1 dimers are very similar, ranging between 38-41°C. In contrast, highly stable SC components SYCE2-TEX12 and SYCP3 have melting temperatures of approximately 65°C (*24, 29*). Thus, the relatively low melting temperatures of SYCE1-SIX6OS1N complexes and SYCE1 suggest that they may undergo conformational change *in vivo*, with each conformation functioning at different stages of meiosis and/or at different locations within the SC. Further, our analysis of SYCE1 infertility-associated mutations and a targeted internal deletion of SIX6OS1 revealed at least four possible conformations of SYCE1 and SYCE1-SIX6OS1 complexes (Figure 7h). Owing to the direct competition between SIX6OS1N-binding and SYCE1core dimerization, these conformations could be achieved in absence of mutations, through alterations of protein levels, local concentrations, allosteric changes and post-translational modifications. As such, alterative conformations of SYCE1 and SYCE1-SIX6OS1 are intriguing candidates for local structural heterogeneity and the propagation of signals along the length of the SC, which could function in roles such as crossover enforcement and interference. Thus, as we progress towards a full molecular understanding of the mammalian SC, the multivalent SYCE1-SIX6OS1 interactions described herein provide tantalising possibilities for a dynamic role of SC structure in its enigmatic functions in the mechanics of meiosis.

## Materials & Methods

### Recombinant protein expression and purification

Human SYCE1 sequences were cloned into pHAT4 and pMAT11 vectors (*45*) for bacterial expression as His- and His-MBP-fusions with TEV cleavage sites for fusion protein removal. Human SIX6OS1 was cloned into pRSF-Duet1 vectors with a TEV-cleavable N-terminal MBP fusion for co-expression with SYCE1. Proteins were expressed in BL21(DE3) *E. coli* cells (Novagen®), in 2xYT media. Expression was induced with addition of 0.5 mM IPTG with the cells incubated at 25 °C for 16 hours. Cells were lysed via sonication in 20 mM Tris pH 8.0, 500 mM KCl, followed by centrifugation. Supernatant was applied to an amylose (NEB) affinity chromatography column, followed by HiTrap Q HP (GE Healthcare) anion exchange chromatography. His- and His-MBP/MBP tags were removed by incubation with TEV protease at 4 °C for 16 hours. The cleaved proteins were further purified by HiTrap Q HP (GE Healthcare) anion exchange chromatography followed by size exclusion chromatography (HiLoad^TM^ 16/600 Superdex 200, GE Healthcare). The purified proteins/complexes were concentrated using Microsep™ Advance 3kDa (PALL) centrifugal filter units and stored at − 80 °C. Protein samples were analysed for purity using Coomassie-stained SDS-PAGE. Protein molecular weights and extinction coefficients were calculated using ExPASY ProtParam (http://web.expasy.org/protparam/) with protein concentrations determined using a Cary 60 UV spectrophotometer (Agilent).

### Circular dichroism (CD)

Far-UV CD spectra were collected using a Jasco J-810 spectropolarimeter (Institute for Cell and Molecular Biosciences, Newcastle University). Wavelength scans were recorded at 4°C from 260 to 185 nm at 0.2 nm intervals using a 0.2 mm pathlength quartz cuvette (Hellma). Protein samples were measured at 0.2-0.4 mg/ml in 10 mM Na_2_HPO_4_ pH 7.5, 150 mM NaF. Nine measurements were taken for each sample, averaged, buffer corrected and converted to mean residue ellipticity ([θ]) (x1000 deg.cm^2^.dmol^-1^.residue^-1^). Spectral deconvolutions were carried out using the Dichroweb CDSSTR algorithm (http://dichroweb.cryst.bbk.ac.uk). CD thermal melts were recorded at 222 nm between 5°C and 95°C, at intervals of 0.5°C with a 1°C per minute ramping rate. Protein samples were measured at 0.1 mg/ml in 20 mM Tris pH 8.0, 150 mM KCl, 2 mM DTT, using a 1 mm pathlength quartz cuvette (Hellma). The data were plotted as % unfolded after conversion to MRE ([θ]_222,x_-[θ]_222,5_)/([θ]_222,95_-[θ]_222,5_). The melting temperature was determined as the temperature at which the proteins are 50% unfolded.

### Size-exclusion chromatography multi-angle light scattering (SEC-MALS)

SEC-MALS analysis of protein samples was carried out at concentrations of 5-20 mg/ml in 20 mM Tris pH 8.0, 150 mM KCl, 2 mM DTT. Samples were loaded onto a Superdex™ 200 Increase 10/300 GL (GE Healthcare) column at 0.5 ml/min using an ÄKTA™ Pure (GE Healthcare) system. The eluate was fed into a DAWN® HELEOS™ II MALS detector (Wyatt Technology), followed by an Optilab® T-rEX™ differential refractometer (Wyatt Technology). SEC-MALS data was collected and analysed using ASTRA® 6 software (Wyatt Technology), using Zimm plot extrapolation with a 0.185 ml/g dn/dc value to determine absolute protein molecular weights.

### Size-exclusion chromatography small-angle X-ray scattering (SEC-SAXS)

SEC-SAXS experiments were carried out on beamline B21 at Diamond Light Source synchrotron facility (Oxfordshire, UK). Protein samples at concentrations 6-20 mg/ml were loaded onto a Superdex™ 200 Increase 10/300 GL size exclusion chromatography column (GE Healthcare) in 20 mM Tris pH 8.0, 150 mM KCl at 0.5 ml/min using an Agilent 1200 HPLC system. The eluate was fed through the experimental cell, with SAXS data recorded at 12.4 keV, in 3.0 s frames with a detector distance of 4.014 m. ScÅtter 3.0 (http://www.bioisis.net) was used to subtract, average the frames and carry out the Guinier analysis for the *Rg* and cross-sectional *Rg* (*Rc*). *P(r)* distributions were fitted using PRIMUS. *Ab initio* modelling was performed using DAMMIF (*46*) imposing P1 symmetry. 20 independent runs were averaged. The PyMOL Molecular Graphics System, Version 2.0 Schrödinger, LLC was used to generate images of the SAXS *ab initio* models.

### Yeast two-hybrid (Y2H)

Constructs of human SYCE1 and SIX6OS1 were cloned into pGBKT7 and pGADT7 vectors (Clontech). Y2H experiments were carried out using the Matchmaker™ Gold system (Clontech) according to manufacturer’s guidelines. Y187 yeast strain was transformed with pGBKT7 vectors while the Y2H gold strain was transformed with pGADT7 vectors. Yeast transformations were carried out using standard lithium acetate methods. Mating of the two strains was carried out in 0.5 ml 2xYPDA at 30°C, 40 r.p.m, by mixing respective colonies. After 24 hours the cultures were centrifuged and pellets were resuspended in 0.5xYPDA. These were then plated onto SD/-Trp/-Leu to select for mated colonies and onto SD/-Trp/-Leu/-Ade/-His with X-α-gal to detect mated colonies through ADE1, HIS3 and MEL1 reporter gene activation. Plates were then incubated for 5 days at 30 °C.

#### Production of CRISPR/Cas9-Edited Mice

For developing the *Syce1^POF/POF^* model, *Syce1*-sgRNA 5’-TGACTTCTTTCCACACTATC-3’ targeting the intron 10 was predicted at https://eu.idtdna.com/site/order/designtool/index/CRISPR_SEQUENCE. This crRNA, the tracrRNA and the ssODN (5’GGGACTCTTCCTCCGAAGCCATGAGGCAGCTGCAGCAATGTAAGATGCAGGGTGGGGCAGGAGGAGGAAATGTCTAGCACTGACTTCTTTCCACACCCCCAGGTAGA TCTTCAAGGATGAGAACAAGAAAGCTGAGGAGTTCCTAGAGGCTGCAGCTCAGCAGCACGAGCAGCTGCAGCAGAGGTGCCACCAGCTACAG-3’) were produced by chemical synthesis at IDT. The ssODN contains the mutated base (C>T, p.Gln241*) and the PAM was mutated by substituting it by the human intron sequence (ACTATCAG > CCCCCAG). The crRNA and tracrRNA were annealed to obtain the mature sgRNA. A mixture containing the sgRNAs, recombinant Cas9 protein (IDT) and the ssODN (30 ng/μl Cas9, 20 ng/μl of each annealed sgRNA and 10 ng/μl ssODN) were microinjected into B6/CBA F2 zygotes (hybrids between strains C57BL/6J and CBA/J) (*47*) at the Transgenic Facility of the University of Salamanca. Edited founders were identified by PCR amplification (Taq polymerase, NZYtech) with primers flanking the exon 11 (Primer F 5’-CTGTAGAGAAACTGATGAAAGT-3’ and R 5’-CAAGAAAATATGAAGAGACATAC - 3’) producing an amplicon of 398 bp for both edited and wild-type alleles, and either direct sequenced or subcloned into pBlueScript (Stratagene) followed by Sanger sequencing, selecting the point mutation in the targeted region of *Syce1* (Supplementary Figure 1). For generating the *Six6os1^Δ10-21/ Δ10-21^* (named as *Six6os1^Δ/ Δ^*), *Six6os1*-crRNA G68 5’-ATCTGTTTGTCAGTTTGGAC – 3’ and *Six6os1*-crRNA G75 5’-TACTTATGTCTTGCTCATAC-3’ targeting the exon 2 and exon 3 and the ssODN (5’-GTTCTTACTTTATGTATGCTCTTTTATATATGGCTTCTGAAAGTTTTATTATTTATTTTACACAGTGTCCAAGATGAATGATAATCTGTTTGTCAGTTTGCAAGACATAAGTATTAAAGAAGATACGATTCAAAGAATTAATAGTAAGTAGTTTTGCATGAAATAAATATTTTAGTCTTTTGGTTTTATCTTATATAGCA–3’) were predicted, produced and microinjected as before. Edited founders with the predicted deletion were identified through PCR using primers flanking this region (Primer F 5’-CACTTACATTTTCCTTTTAAGAATGC-3’ and R 5’-CCCCTCTCATACATACAAGTTGC-3’). The Δ10-21 allele was 285 bp long versus 413 bp of the wild-type allele (Supplementary Figures 5a and 5b). The founders were crossed with wild-type C57BL/6J to eliminate possible unwanted off-targets. Heterozygous mice were re-sequenced and crossed to give rise to edited homozygous. Genotyping was performed by analysis of the PCR products of genomic DNA with primers F and R.

#### Histology

For histological analysis of ovaries, after the necropsy of the mice their ovaries were removed and fixed in formol 10%. They were processed into serial paraffin sections and stained with haematoxylin-eosin. The samples were analysed using a microscope OLYMPUS BX51 and images were taken with a digital camera OLYMPUS DP70.

#### Immunocytology

Testes were detunicated and processed for spreading using a conventional ‘dry-down’ technique. Oocytes from fetal ovaries (E17.5 embryos) were digested with collagenase, incubated in hypotonic buffer, disaggregated and fixed in paraformaldehyde. Both meiocytes preparations were incubated with the following primary antibodies for immunofluorescence: rabbit αSIX6OS1 R1 and R2 (1:100, Proteogenix (*11*)), rabbit αSYCE1 17406-1-AP (1:50, Proteintech), guinea pig αSYCE1 (1:100, provided by Dr. C. Höög), mouse αSYCP3 IgG sc-74569 (1:1000, Santa Cruz), rabbit serum αSYCP3 K921 (1:500), rabbit αSYCP1 IgG ab15090 (1:200), guinea pig αSYCE3(1:20, provided by Dr R. Benavente), guinea pig αSYCE2 (1:100, provided by Dr. C. Höög), rabbit αTEX12 IgG (1:100, provided by Dr R. Benavente), rabbit anti-γH2AX (ser139) IgG #07-164 (1:200) (Millipore), mouse αMLH1 51-1327GR (1:5, BD Biosciences), rabbit αRAD51 PC130 (1:50, Calbiochem) and rabbit αDMC1 R1 and R2 (1:500, Proteogenix). The secondary antibodies used were goat Alexa 555 α-mouse A-32727, goat Alexa 488 α-mouse A-11001, donkey Alexa 555 α-rabbit A-31572 (1:200, ThermoFisher), goat Alexa 488 - Fab α-rabbit 111-547-003 and donkey FITC α-guinea pig 706-095-148 (1:100, Jackson Immunoresearch). Slides were visualized at room temperature using a microscope (Axioplan 2; Carl Zeiss, Inc.) with 63 × objectives with an aperture of 1.4 (Carl Zeiss, Inc.). Images were taken with a digital camera (ORCA-ER; Hamamatsu) and processed with OPENLAB 4.0.3 and Photoshop (Adobe). Quantification of fluorescence signals was performed using ImageJ software.

#### Cell lines and transfections

HEK 293T and COS7 cell lines were and obtained from the ATCC. Cell lines were tested for mycoplasma contamination (Mycoplasma PCR ELISA, Sigma). They were transfected with Jetpei (PolyPlus) according to the manufacturer protocol.

#### Immunoprecipitation and western blotting

HEK293T cells were transiently transfected and whole cell extracts were prepared and cleared with protein G Sepharose beads (GE Healthcare) for 1 h. The antibody was added for 2 h and immunocomplexes were isolated by adsorption to protein G-Sepharose beads o/n. After washing, the proteins were eluted from the beads with 2xSDS gel-loading buffer 100mM Tris-Hcl (pH 7), 4% SDS, 0.2% bromophenol blue, 200mM β-mercaptoethanol and 20% glycerol, and loaded onto reducing polyacrylamide SDS gels. The proteins were detected by western blotting with the indicated antibodies. Immunoprecipitations were performed using mouse αFlag IgG (5µg; F1804, Sigma-Aldrich), mouse αGFP IgG (4 µg; CSB-MA000051M0m, Cusabio), mouse αMyc obtained from hybridoma cell myc-1-9E10.2 ATCC (4 µg) and ChromPure mouse IgG (5µg/1mg prot; 015-000-003). Primary antibodies used for western blotting were rabbit αFlag IgG (1:2000; F7425 Sigma-Aldrich), goat αGFP IgG (sc-5385, Santa Cruz) (1:3000), rabbit αMyc Tag IgG (1:3000; #06-549, Millipore). Secondary horseradish peroxidase-conjugated α-mouse (715-035-150, Jackson ImmunoResearch), α-rabbit (711-035-152, Jackson ImmunoResearch), or α-goat (705-035-147, Jackson ImmunoResearch) antibodies were used at 1:5000 dilution. Antibodies were detected by using Immobilon Western Chemiluminescent HRP Substrate from Millipore. Both *Syce1*POF and *Six6os1* Δ10-21 cDNAs used for IF and co-IP experiments were RT-PCR amplified (the primers used for it were Syce1 S 5’-GAGCAGTATGGCCACCAGACC-3’ and Syce AS 5’-GAGGAGGGTATTAGGTCCTGC-3’; Six6os1 S 5’-AGTGTCCAAGATGAATGATAATCTG-3’ and Six6os1 AS 5’-GTTCAAAAATAATAACTCAAAAAAAC-3’) from total RNA extracted from *Syce1^POF/POF^* and *Six6os1^Δ10-21/ Δ10-21^* mice respectively. PCR amplified fragments were cloned in pcDNA3-based mammalian expression vectors with different tags (EFGP or Flag) and verified by Sanger sequencing.

#### Statistics

In order to compare counts between genotypes, we used the Welch’s t-test (unequal variances t-test), which was appropriate as the count data were not highly skewed (i.e., were reasonably approximated by a normal distribution) and in most cases showed unequal variance. We applied a two-sided test in all the cases. Asterisks denote statistical significance: *p-value <0.01, **p-value <0.001 and ***p-value<0.0001.

#### Ethics statement

Mice were housed in a temperature-controlled facility (specific pathogen free, spf) using individually ventilated cages, standard diet and a 12 h light/dark cycle, according to EU laws at the “Servicio de Experimentación Animal, SEA”. Mouse protocols were approved by the Ethics Committee for Animal Experimentation of the University of Salamanca (USAL). We made every effort to minimize suffering and to improve animal welfare. Blinded experiments were not possible since the phenotype was obvious between wild type and mutant mice for all of the experimental procedures used. No randomization methods were applied since the animals were not divided in groups/treatments. The minimum size used for each analysis was two animals/genotype.

## Acknowledgements

We thank Diamond Light Source and the staff of beamline B21 (proposals sm15836, sm21777 and sm23510). We thank H. Waller for assistance with CD data collection. ORD is a Sir Henry Dale Fellow jointly funded by the Wellcome Trust and Royal Society (Grant Number 104158/Z/14/Z). This work was supported by MINECO (BFU2017-89408-R) and by Junta de Castilla y Leon (CSI239P18). FSS, LGH and NFM are supported by European Social Fund/JCyLe grants (EDU/556/2019, EDU/1083/2013 and EDU/310/2015). CIC-IBMCC is supported by the Programa de Apoyo a Planes Estratégicos de Investigación de Estructuras de Investigación de Excelencia cofunded by the Castilla–León autonomous government and the European Regional Development Fund (CLC–2017–01). The funders had no role in study design, data collection and analysis, decision to publish, or preparation of the manuscript.

## Author contributions for biochemical data

FSS, OMD, LGH, NFM, CG-P, MSM and ORD. performed experiments. ORD and AMP designed experiments, analysed data and wrote the manuscript together with AMP. AMP, ELC and ORD supervised and designed the work.

## Supplementary Materials

**Supplementary Figure 1.**
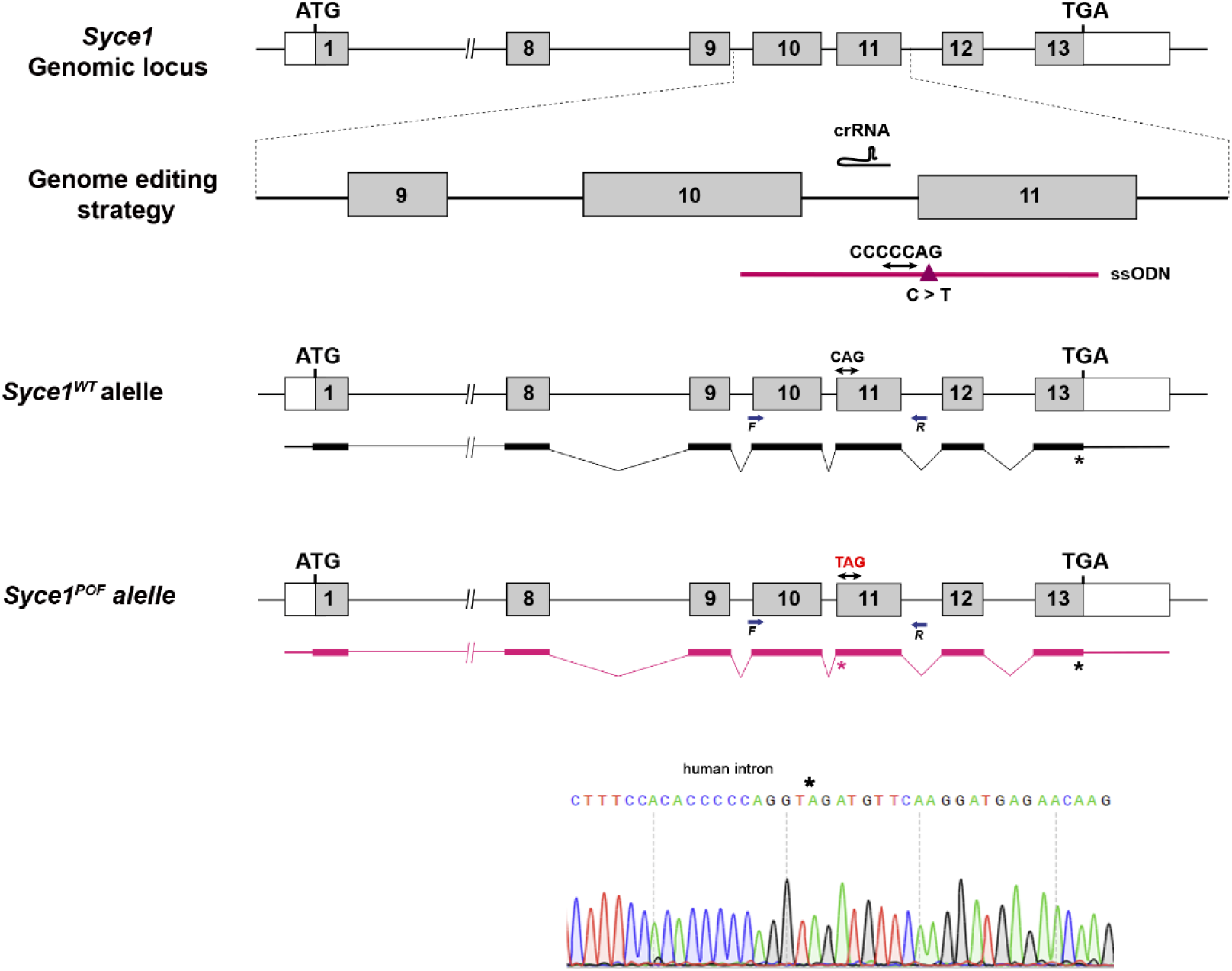
Generation of *Syce1^POF/POF^* mouse model. Diagrammatic representation of the mouse *Syce1* locus (WT) and the genome editing strategy showing the crRNA located on intron 10 and the ssODN targeting a region between exon 10 and 11 carrying the point mutation (c.613C>T, p.Gln241*) in addition to the PAM substituted by the human intron sequence (see methods). The corresponding coding exons (grey boxes) and non-coding exons (open boxes) are represented. Thin (non-coding) and thick (coding sequences) lines under exons represent the expected transcript derived from WT (black) and *Syce1POF* edited allele (pink). ATG, initiation codon; TGA and *, stop codon. The newly generated stop codon in the edited transcript is indicated (*pink). Primers for PCR genotyping (F and R) are represented by arrows. The nucleotide sequence of the edited region derived from PCR amplification of DNA from the *Syce1^POF/POF^* is indicated.

**Supplementary Figure 2.**
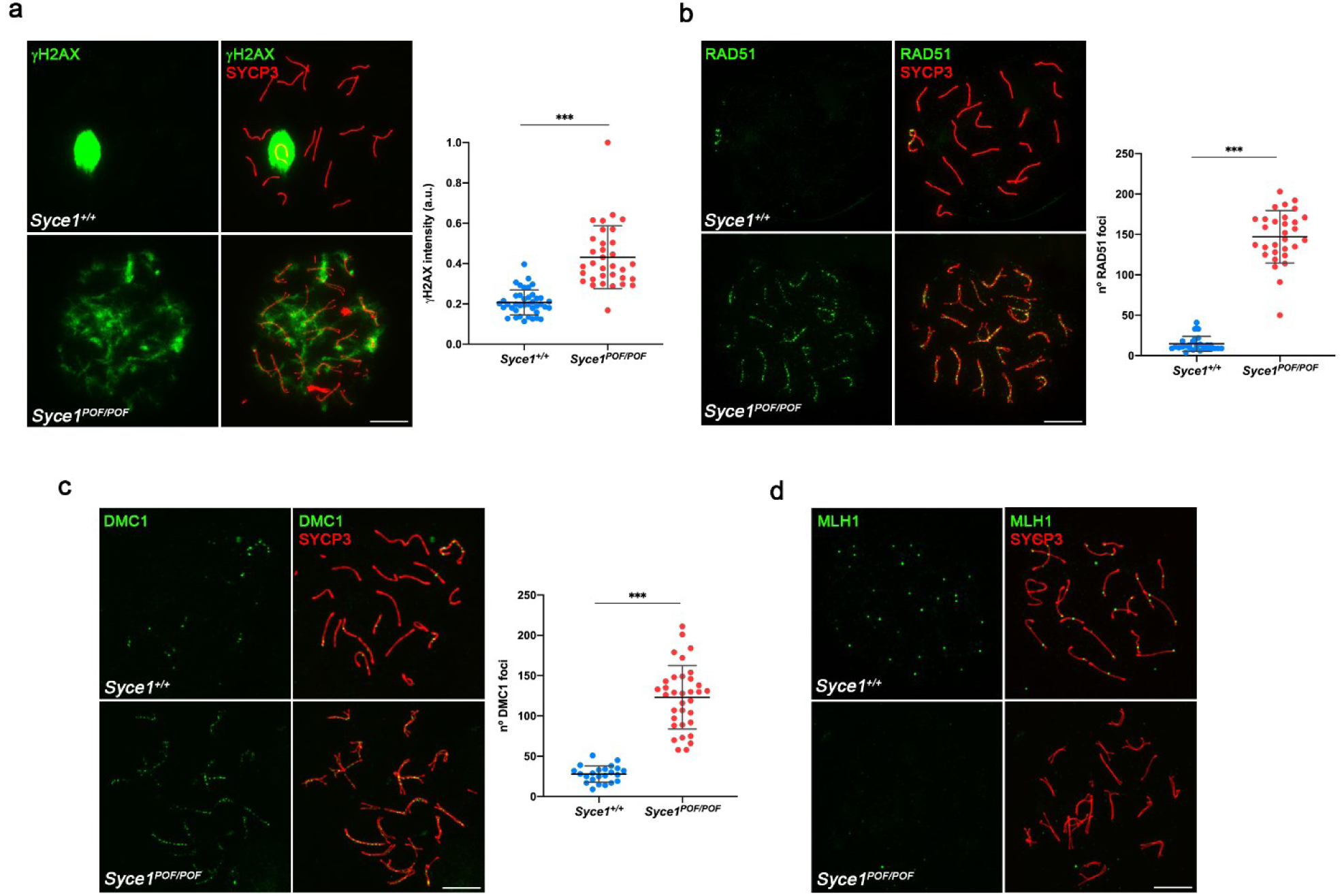
DSBs are deficiently repaired in *Syce1^POF/POF^* spermatocytes. **(a)** Double immunolabeling of γ-H2AX (green) and SYCP3 (red) in spermatocyte spreads from WT and *Syce1^POF/POF^* mice. γ-H2AX persists labeling the chromatin of *Syce1^POF/POF^* pachytene-like in contrast to the WT pachytenes where the labeling is restricted to the sex body. **(b-c)** Double immunofluorescence of **(b)** RAD51 or **(c)** DMC1 (green) and SYCP3 (red). *Syce1^POF/POF^* pachytene-like spermatocytes show increased number of foci of both RAD51 and DMC1 along the AEs compared to the WT indicating unrepaired DSBs. **(d)** Double immunolabeling of MLH1 (green) and SYCP3 (red) showing the absence of COs (MLH1) in the arrested *Syce1^POF/POF^* spermatocytes. Scale bars represent 10 μm. Plots right to the image panels represent the quantification of fluorescence intensity **(a)** or number of foci **(b-c)** from WT and pachytene-like arrested spermatocytes. Welch’s *t*-test analysis: * p<0.01, **p<0.001, ***p<0.0001.

**Supplementary Figure 3.**
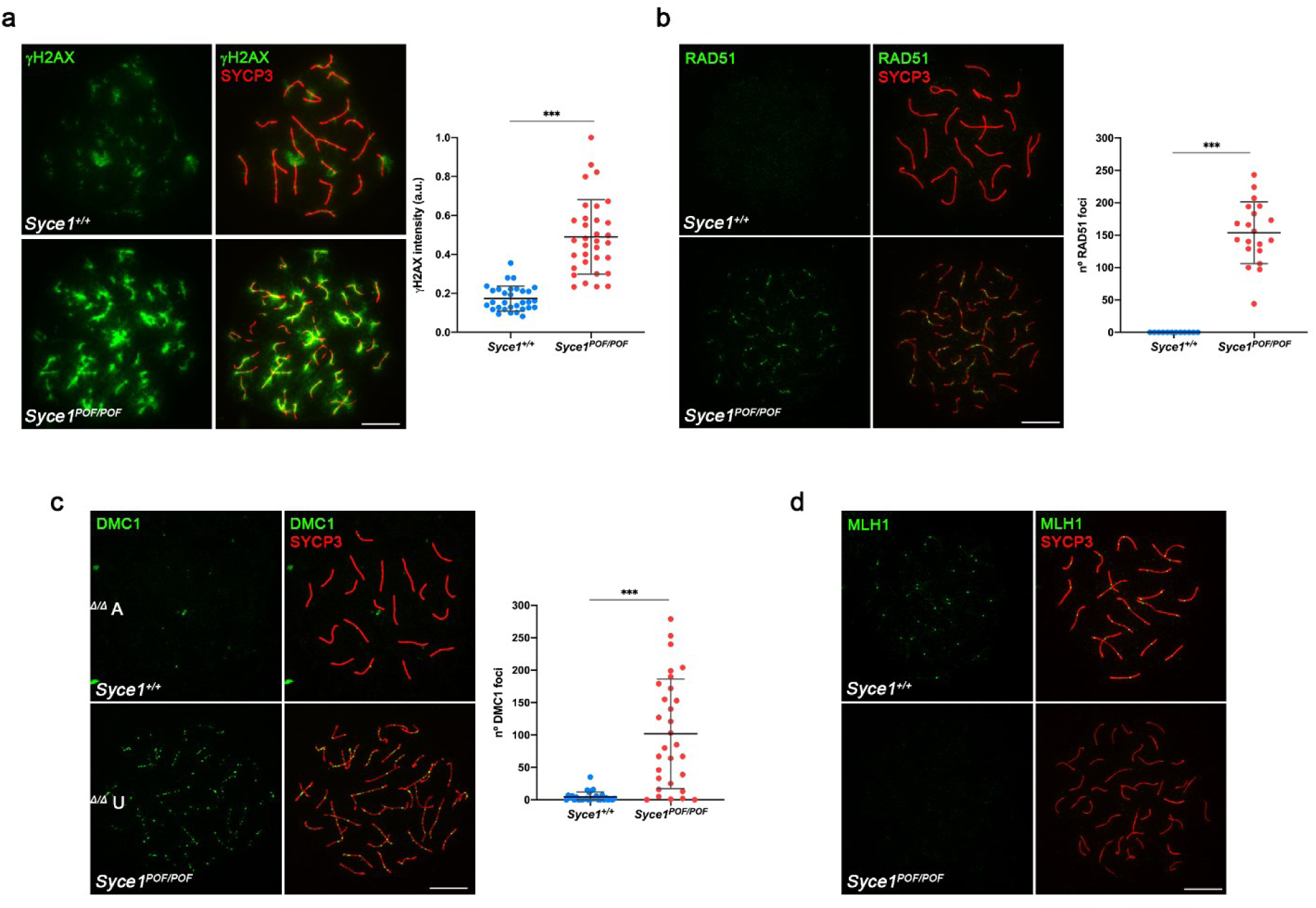
*Syce1^POF/POF^* oocytes do not repair properly DSBs. **(a)** Double immunostaining of spread preparations of WT pachytene and *Syce1^POF/POF^* pachytene-like oocytes with γ-H2AX (green) and SYCP3 (red). In *Syce1^POF/POF^* oocytes the levels of γ-H2AX are increased and more restricted to the AEs compared to the WT pachytenes. **(b-c)** Double immunolabeling of **(b)** RAD51 or **(c)** DMC1 (green) and SYCP3 (red), showing higher number of foci in the AEs from mutant oocytes. **(d)** Labeling of MLH1 (green) and SYCP3 (red). MLH1 foci are absent at the AEs of *Syce1^POF/POF^* oocytes. Scale bars represent 10 μm. Plots right to the image panels represent the quantification of fluorescence intensity **(a)** or number of foci **(b-c)** from WT and pachytene-like arrested oocytes. Welch’s *t*-test analysis: * p<0.01, **p<0.001, ***p<0.0001.

**Supplementary Figure 4.**
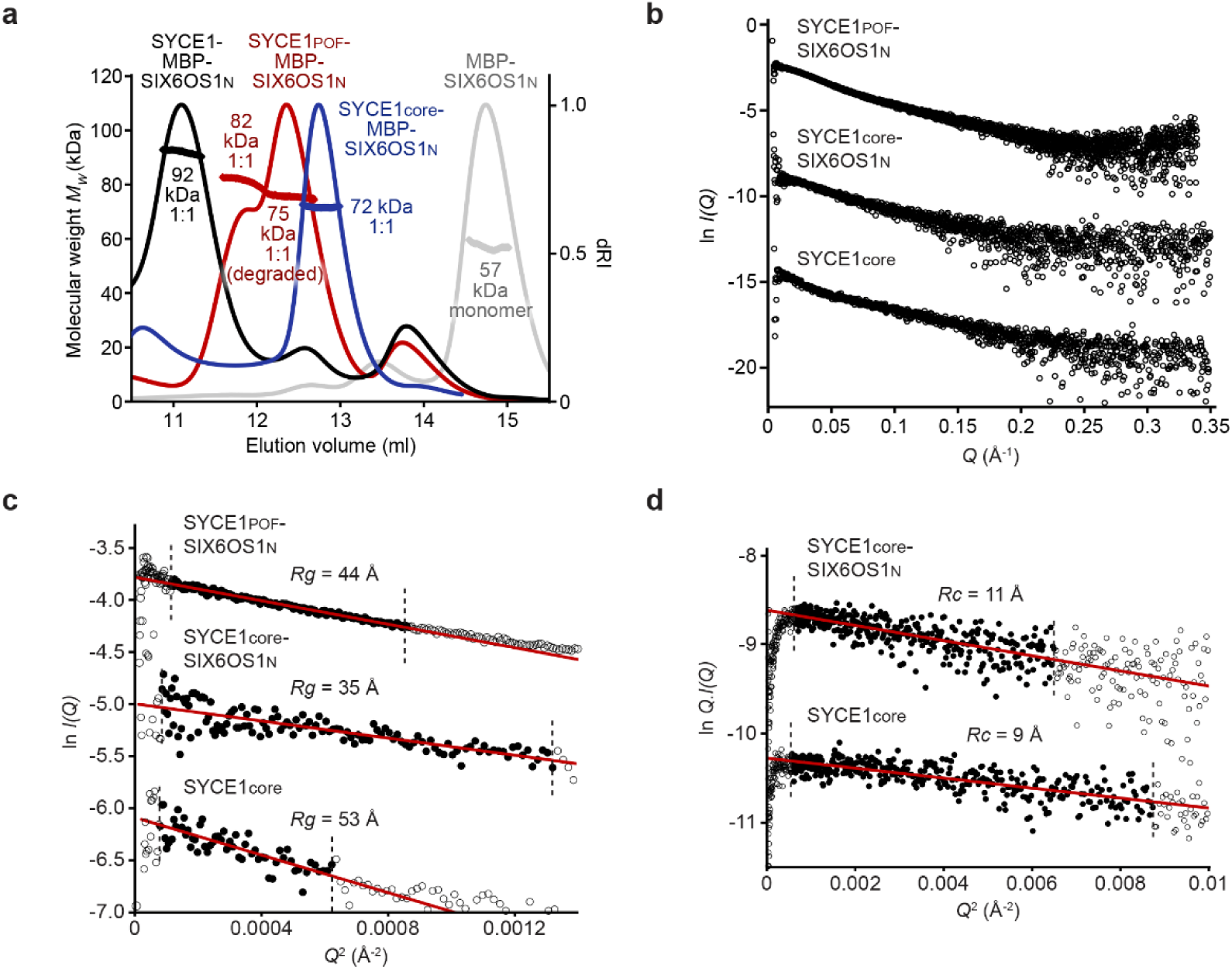
SEC-MALS and SEC-SAXS analysis. (**a**) SEC-MALS analysis of His- and MBP-tagged SYCE1-SIX6OS1 complexes. SYCE1core-SIX6OS1N (blue), SYCE1POF-SIX6OS1N (red) and full-length SYCE1-SIX6OS1N (black) are 1:1 complexes of 72 kDa, 82 kDa (75 kDa for the degradation product complex) and 92 kDa, respectively (theoretical 1:1 – 75 kDa, 83 kDa and 95 kDa). (**b-d**) SEC-SAXS analysis of SYCE1core-SIX6OS1N, SYCE1POF-SIX6OS1N and SYCE1core. (**b**) SEC-SAXS scattering curves. (**c**) SEC-SAXS Guinier analysis to determine the radius of gyration (*Rg*). The linear fits are highlighted in black and are demarcated by dashed lines. The *Q*.*Rg* values were < 1.3. (**d**) SEC-SAXS Guinier analysis to determine the radius of gyration of the cross-section (*Rc*). The linear fits are highlighted in black and are demarcated by dashed lines. The *Q*.*Rc* values were < 1.3.

**Supplementary Figure 5.**
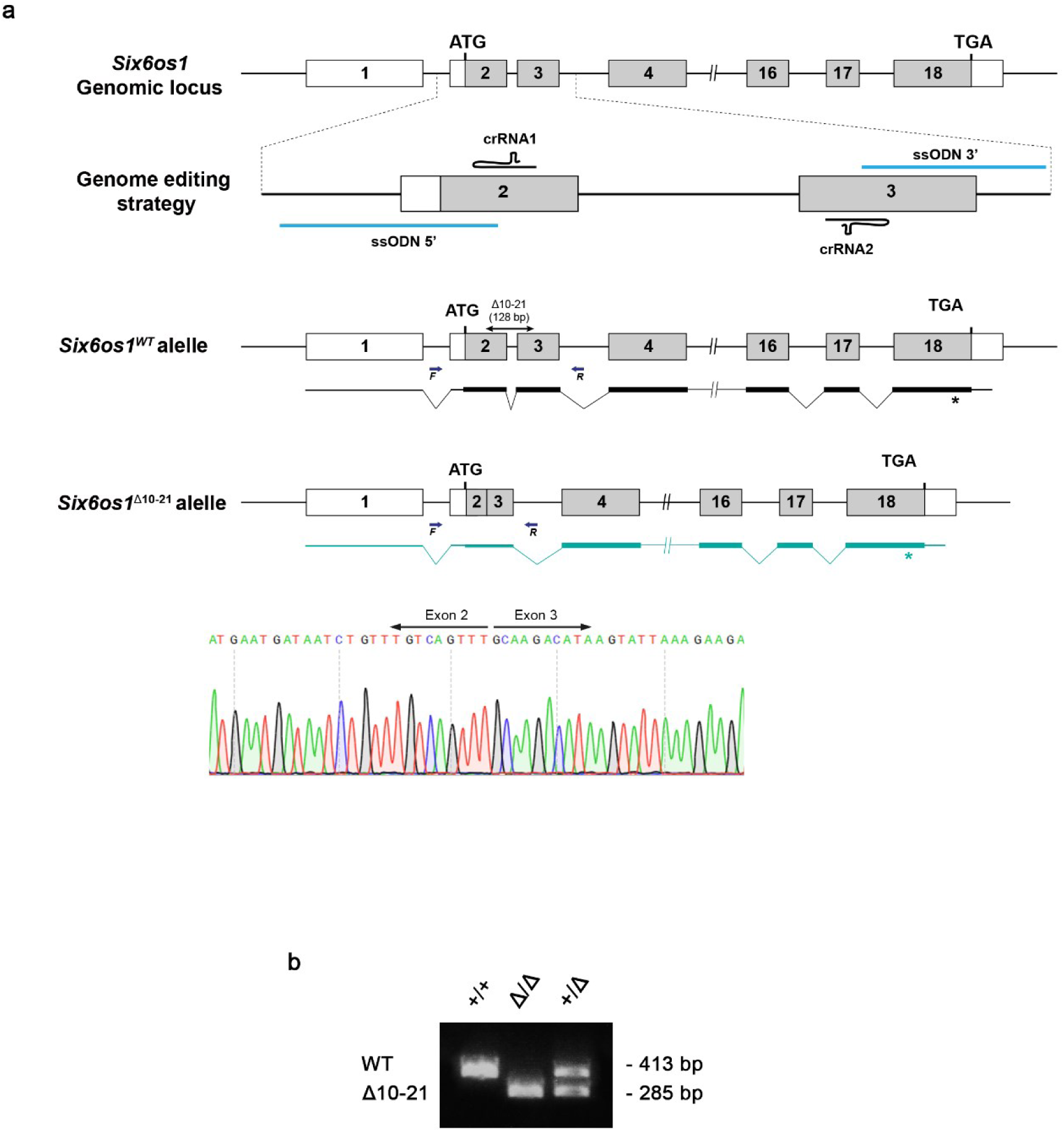
Generation of *Six6os1*^Δ^*^10-21/^*^Δ^*^10-21^* mutant mice. **(a)** Schematic representation of the genome editing strategy for the generation of the *Six6os1* Δ10-21 mutant mouse model showing the crRNAs located on exons 2 and 3. The ssODN comprises the ssODN 5’ and the ssODN 3’ sequences, targeting the 5’ of exon 2 and the 3’ of exon 3 respectively (see methods). The corresponding coding exons (grey boxes) and non-coding exons (open boxes) are represented. Thin (non-coding) and thick (coding sequences) lines under exons represent the expected transcript derived from wild-type (black) and *Six6os1* Δ10-21 edited allele (green). ATG, initiation codon; TGA and *, stop codon. Primers for PCR genotyping (F and R) are represented by arrows. The nucleotide sequence of the edited region showing the deletion of 128 bp derived from PCR amplification of DNA from the *Six6os1*^Δ*10-21/*Δ*10-21*^ (named in the figures *Six6os1*^Δ*/*Δ^). **(b)** PCR analysis of genomic DNA from three littermate progeny of *Six6os1*^Δ*10-21/WT*^ crosses. The PCR amplification with primers F and R revealed 413 and 285 bp fragments for wild-type and edited alleles respectively. Wild-type (+/+), Δ10-21 homozygous (Δ/Δ) and heterozygous (Δ/+) animals.

**Supplementary Figure 6.**
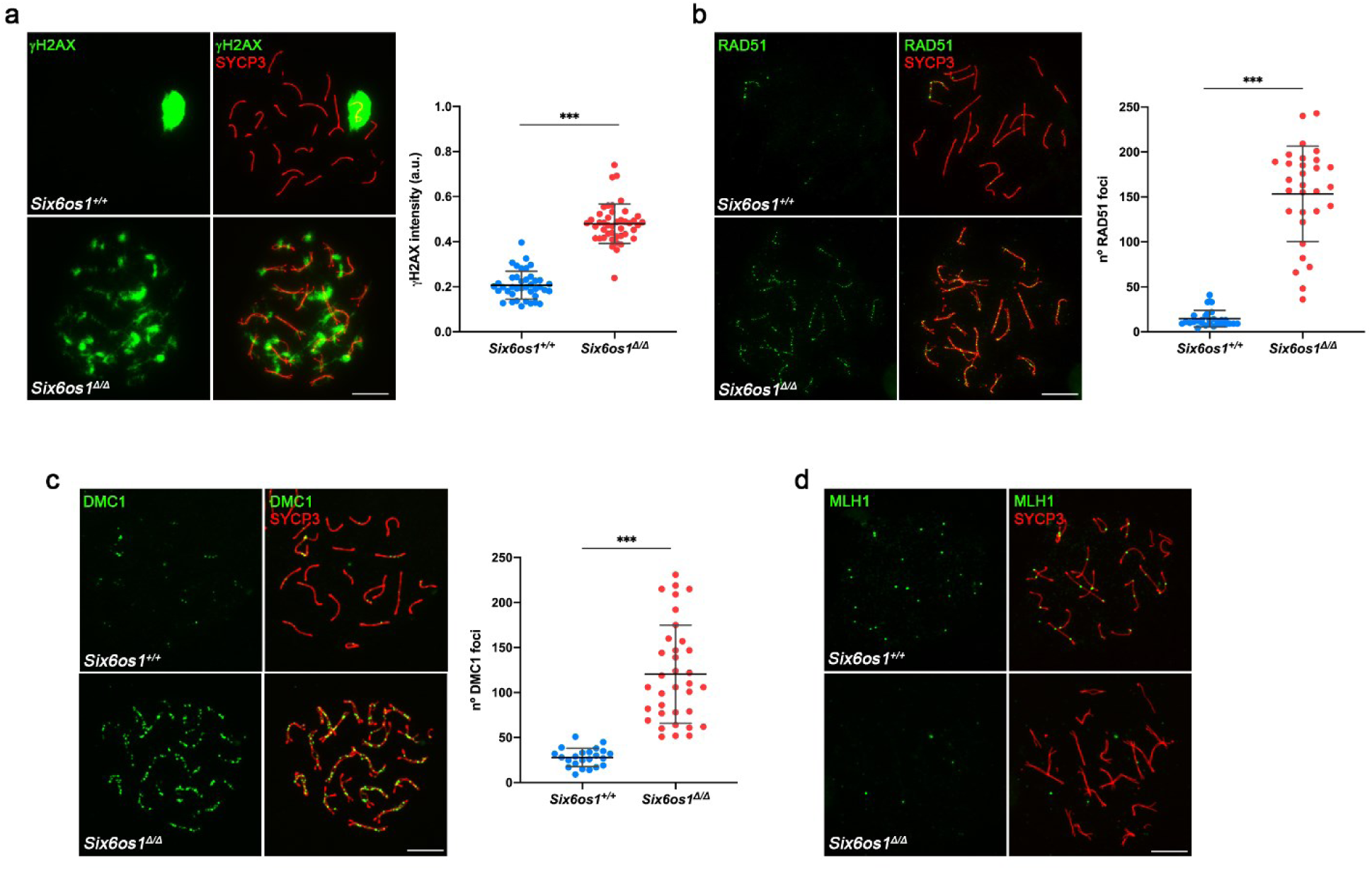
DSBs are generated but defectively repaired in *Six6os1*^Δ*10-21*/Δ^*^10-21^* spermatocytes. **(a)** Double immunofluorescence of γ-H2AX (green) and SYCP3 (red) in spermatocyte spreads from WT and *Six6os1*^Δ*/*Δ^ mice. In WT pachytenes γ-H2AX intensely labels the chromatin of the sex bivalent, while in the *Six6os1*^Δ*/*Δ^ arrested pachytene-like γ-H2AX labeling remains in the chromatin. **(b-c)** Double immunolabeling of **(b)** RAD51 or **(c)** DMC1 (green) and SYCP3 (red). Both RAD51 and DMC1 remain associated to the AEs in the *Six6os1*^Δ*/*Δ^ pachytene-like spermatocytes, showing increased number of foci than the WT. **(d)** Double immunolabeling of MLH1 (green) and SYCP3 (red). MLH1 is absent from the AEs in the *Six6os1*^Δ*/*Δ^ arrested spermatocytes. Scale bars represent 10 μm. Plots under the image panels represent the quantification of fluorescence intensity **(a)** or number of foci **(b-c)** from WT and pachytene-like arrested spermatocytes. Welch’s *t*-test analysis: * p<0.01, **p<0.001, ***p<0.0001.

